# Sex chromosome gene expression associated with vocal learning following hormonal manipulation in female zebra finches

**DOI:** 10.1101/2021.07.12.452102

**Authors:** Matthew H. Davenport, Ha Na Choe, Hiroaki Matsunami, Erich D. Jarvis

## Abstract

Zebra finches are sexually dimorphic vocal learners. Males learn to sing by imitating mature conspecifics, but females do not. Absence of song in females is associated with partial atrophy and apparent repression of several vocal learning brain regions during development. However, atrophy can be prevented and vocal learning retained in females when given early pharmacological estrogen treatment. To screen for candidate drivers of this sexual dimorphism, we performed an unbiased transcriptomic analysis of song learning nuclei specializations relative to the surrounding regions from either sex, treated with vehicle or estrogen until 30 days old when divergence between the sexes becomes anatomically apparent. Analyses of transcriptomes by RNA sequencing identified song nuclei-specialized gene expressed modules associated with sex and estrogen manipulation. Female HVC and Area X gene modules were specialized by estrogen supplementation, exhibiting a subset of the transcriptomic specializations observed in males. Female RA and LMAN specialized modules were less dependent on estrogen. The estrogen-induced gene modules in females were enriched for anatomical development functions and strongly correlated to the expression of several Z sex chromosome genes. We present a hypothesis where reduced dosage and expression of these Z chromosome genes suppresses the full development of the song system and thus song learning behavior, which is partially rescued by estrogen treatment.

## Introduction

Vocal learning is the ability to imitate heard sounds using a vocal organ, and is a necessary and specialized component for spoken language and song. Vocal learning is found in seven non-human clades, four mammalian and three avian, each having independently evolved the trait^1,2^. Oscine songbirds have proved to be the most tractable for studying vocal learning in the lab, with much of the field focusing on the Australian zebra finch (*Taeniopygia castanotis*). Despite ∼300 million years of separation from their common ancestor^3,4^, there is remarkable evolutionary convergence between songbird and human vocal learning in terms of behavioral progression, developmental effects of deafening, anatomical connectivity of vocal-motor learning circuits, sites of accelerated evolution within the genome, and genes with specialized up- or down-regulated expression in song and speech circuits relative to the surrounding motor control circuits^1,5–13^. Unlike in humans, however, vocal learning is strongly sexually dimorphic in zebra finches and many other vocal learning species^14,15^. Male zebra finches learn to produce a species appropriate song by imitating mature male conspecifics during juvenile development, while females are limited to producing innate calls^16^.

The songbird vocal motor learning circuit contains four major interconnected telencephalic song control nuclei; HVC (proper name) in the dorsal nidopallium (DN); the lateral magnocellular nucleus of the anterior nidopallium (LMAN) in the anterior nidopallium (AN); the robust nucleus of the arcopallium (RA) in the lateral intermediate arcopallium (LAI; also called AId); and Area X in the striatum (Str; **Fig. 1a**)^7^. During juvenile development in zebra finch females, HVC and RA atrophy, HVC fails to form synapses in RA, and Area X never appears^14,17–24^. Amazingly, female zebra finches treated with estrogen or a synthetic analog at an early age do not exhibit song system atrophy and instead form a functional neural circuit with all the anatomical components and connections seen in males^25–28^. This “masculinized” song system allows estrogen supplemented females to imitate vocalizations, though not with the same accuracy as males^25,27,29–31^. Interestingly, lesioning female HVC prevents estrogen dependent anatomical “masculinization” of its postsynaptic targets to RA and Area X^32^.

**Figure 1 -.**
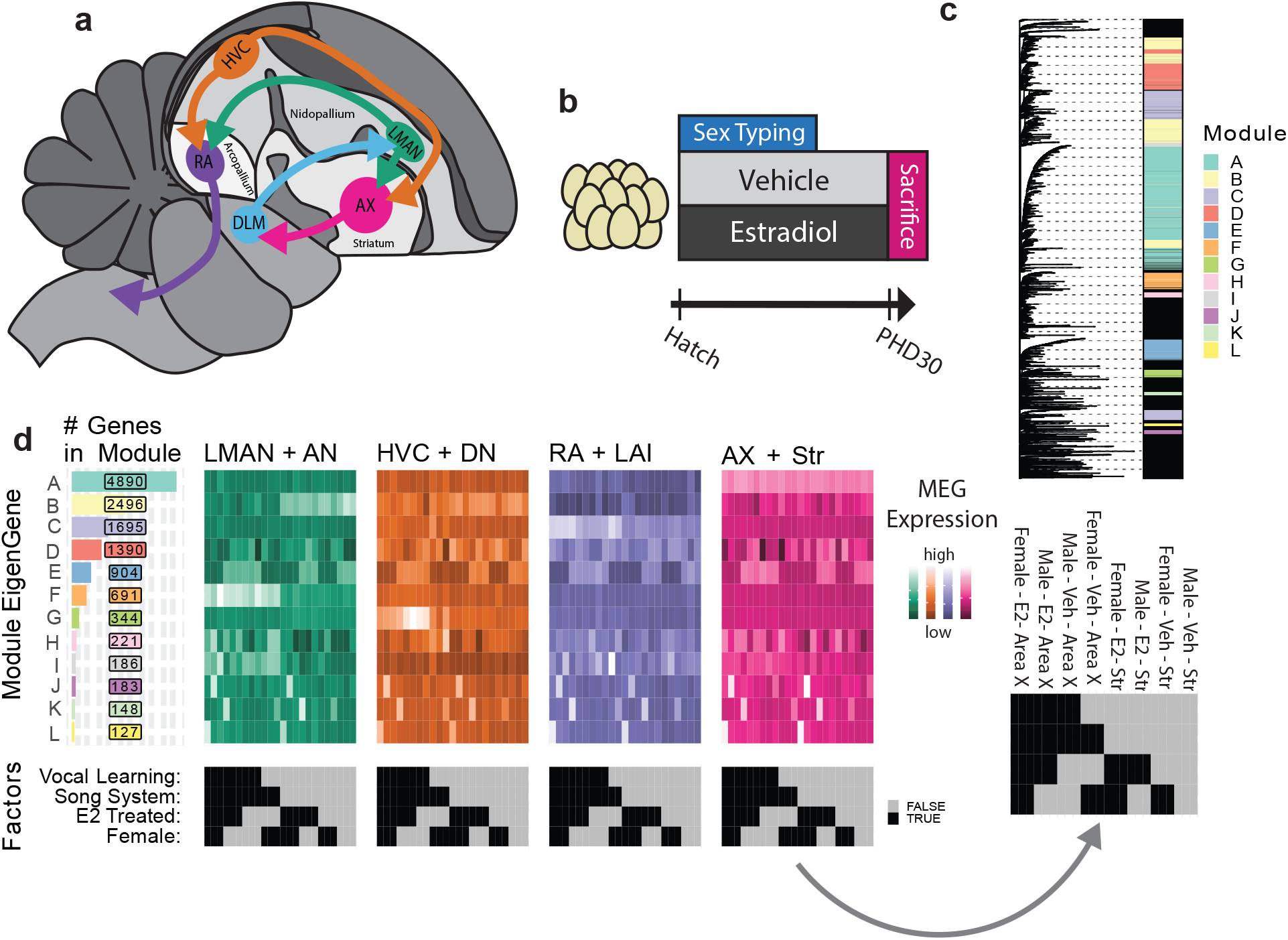
Song system anatomy and experimental design. **a,** Diagram of song system connectivity within the adult male zebra finch brain with major telencephalic domains indicat- ed. Area X connects back to LMAN through the non-vocal specific thalamic nucleus DLM. **b,** Experimental design. Animals were treated with E2 or a vehicle from hatch until sacrifice on PHD30. **c,** WGCNA assignment of genes to modules. Left: Hierarchy computed over the transcriptome wide topological overlap matrix of gene to gene correlations in transcript abundance across samples. Right: Module assignment raster, rows are genes colored according to the assigned module, unassigned genes in black. **d,** MEG expression heat- maps arranged by module size (left) aligned to traits of interest (bottom). Each row is an MEG and each sample is a column. Samples are grouped according to neural circuit node in different colored subpanels. Color intensity encodes MEG expression as calculated by WGCNA. An example raster with sample category labels is provided at right.

The genetic basis of this estrogen-sensitive and sexually dimorphic vocal learning in zebra finch remains largely unknown beyond the downstream recruitment of the androgen receptor (AR)^33^. However, the examination of a rare gynandromorphic zebra finch with lateralized sex chromosome composition indicates that genetically male HVC and Area X are larger than their female analogs independent of gonadal hormone production, implicating sex chromosome gene expression within the song system^34^. Unlike the mammalian X and Y, birds have a Z and W sex determination system where females are hemizygous (ZW) and males homozygous (ZZ)^35^. The relevant transcriptional machinery appears to be set up by post-hatch day 30 (PHD30), after which estrogen fails to masculinize female song nuclei or behavior and the male song system enlarges while the female song system atrophies^17,19,25,36^. Taken together with the hypothesis that vocal learning in females was lost multiple independent times amongst songbirds^15^, the extant findings suggest genetic drivers of vocal learning loss associated with estrogen, sex chromosomes, and song nuclei gene expression specializations. To screen for potential genetic drivers (loci whose expression/inheritance regulate the trait) and locate their action within the song system, we performed an unbiased analysis of transcriptomes from song nuclei and surrounding motor control regions in zebra finches of either sex chronically treated with 17-β-estradiol (E2) or vehicle from hatch until sacrifice at PHD30 (**Fig. 1b**). While the birds for this study were sacrificed prior to the developmental presentation of song behavior, we have previously shown that female finches treated with E2 in the same exact way go on to produce rudimentary imitative songs as adults, consistent with the known induction of vocal learning in females by E2^25,29^. We used a new zebra finch genome assembly and annotation produced by the Vertebrate Genomes Project containing both the Z and W chromosomes^37^.

## Results

### Identification of gene expression modules

We first sought to characterize differences in song nuclei gene expression specializations relative to their immediate surrounding motor brain regions in juvenile males and females with and without E2 treatment. We used RNA-Seq data from a previous study by our lab, on the effects of E2 manipulation on the song system in both males and females^29^. This E2-treatment program masculinized females sufficiently for them to produce song as adults in a parallel cohort of birds^29^. Birds were sacrificed as juveniles at post-hatch day 30 (PHD30), after a one hour period of silence to limit activity-dependent gene expression in song nuclei and surrounding motor regions; this developmental age was chosen because it is around the time when both males and E2-treated females start to sing (e.g. subsong) and all four song nuclei are sufficiently developed to be visually apparent in histological sections^29^. The four major song nuclei (HVC, LMAN, RA, and Area X) and their adjacent surrounding motor regions (DN, AN, LAI, and Str respectively; **Fig. 1a**) were dissected using laser capture. In the case of vehicle-treated females, which lack Area X, a piece of striatum was taken from where Area X would be in males to serve as the Area X sample. In our previous study, we found that these PHD30 vehicle-treated males had larger RAs and HVCs than their female counterparts, Area X was absent in vehicle-treated females, and LMAN was similarly sized in both sexes. Following E2 treatment, E2-treated male RA was significantly smaller than vehicle-treated males; while Area X appeared in E2-treated females and HVC was significantly larger than in vehicle-treated females^29^. This past study mapped the RNA-Seq reads to an older genome assembly lacking the W sex chromosome^29^.

With this data, we first remapped RNA-Seq reads (from n = 3 birds per group) to a new zebra finch assembly (GCF_008822105.2) with both the Z and W chromosomes. As many W chromosome genes are duplicated from the Z chromosome and thus highly similar in sequence, only single-mapped reads were considered to minimize misattributed Z chromosome reads. In our quality control analyses, hierarchical clustering of sample expression vectors revealed two technical outliers, one HVC from a vehicle-treated male and one RA from a E2-treated female, which we excluded from further analysis (**Fig. S1**). Similar to previous PCA and hierarchical clustering^29^, the remapped RNA-Seq data expression levels with outliers removed still resulted in separation of song nucleus and surround and some E2-treated female samples, without sufficient resolution at finer group levels (**Fig. S1**). To address our questions at finer resolution and in an unbiased way, we performed weighted gene correlation network analysis (WGCNA), where we decomposed the transcriptome into actively expressed gene modules using hierarchical clustering of the gene adjacency matrix. This matrix describes the inferred structure of gene networks within our data and was calculated with a soft thresholding of the matrix of gene-to-gene expression correlations across samples (**Fig. S2**)^38^. After the genes were given initial hierarchical cluster-based module assignment, they were then iteratively reassigned to the module whose aggregate expression they best correlated with until no additional genes met the WGCNA reassignment threshold (**Fig. S3** and **Methods** for parameterization). This reassignment was performed to ensure that genes are matched to the aggregate measure that best represents their expression.

Of the ∼21,000 annotated zebra finch genes, 13,220 were well expressed in the finch telencephalic brain regions sampled, a comparable number of genes to what we have seen expressed in adult zebra finch telencephalon^39,40^. These 13,220 were assigned to 14 co-expression modules (**Fig. S4a**). The median observation of unassigned transcripts was 22-fold lower than the median observation of module assigned transcripts (4.86 vs. 0.22 FPKM). Two modules clearly marked single samples from two different birds (**Fig. S4a,b - arrows**), indicative of technical overfitting; these two modules (not birds) were excluded from further analysis, resulting in a more visibly diverse pattern within and across modules (**Fig. S4c,d**). The remaining 12 modules contained 12,444 genes in total; these 12 moduleswere lettered in descending order of size A through L, containing from 4,890 to 127 constituent genes each, which were dynamically expressed across brain regions and treatment groups (**Fig. 1c**). The results of module assignment can be found in supplemental_gene_module_assignment.csv.

### Song nuclei specialization modules and sex differences

To understand song system transcriptional specialization in the context of the gene modules, we calculated the module eigengene (MEG) expression values for each of the 12 modules (**Fig. 1d**). MEGs are the first principle component of variance of all genes in a module and are the aggregate measure for each module’s expression across samples. We then tested for statistically significant correlations between MEG expression for each module and the song nuclei specializations relative to their respective surrounds. Each song nucleus in vehicle-treated control males had unique and overlapping specialization of genes in 2 to 4 modules of the 12 (**Fig. 2a**). The male LMAN specialization relative to the surrounding AN were correlated with modules B, F and I; male HVC specialization relative to DN consisted of genes in modules B, F, and G; male RA specialization relative to LAI consisted of genes in modules C and L; and male Area X specialization relative to the surrounding Str consisted of genes in modules C, F, G, and I. Area X was the only song nucleus that did not have a specialized module unique to it relative to other song nuclei (**Fig. 2a**). In non-vocal learning juvenile females, interestingly LMAN was specialized relative to AN by the same gene modules as in males (B, F, and I) as well as an additional module G (**Fig. 2b**); RA was specialized by module A as in males, but not module L and by additional modules A and G. In contrast, neither juvenile female HVC nor Area X exhibited significant gene module expression specializations relative to their surrounds.

**Figure 2 -.**
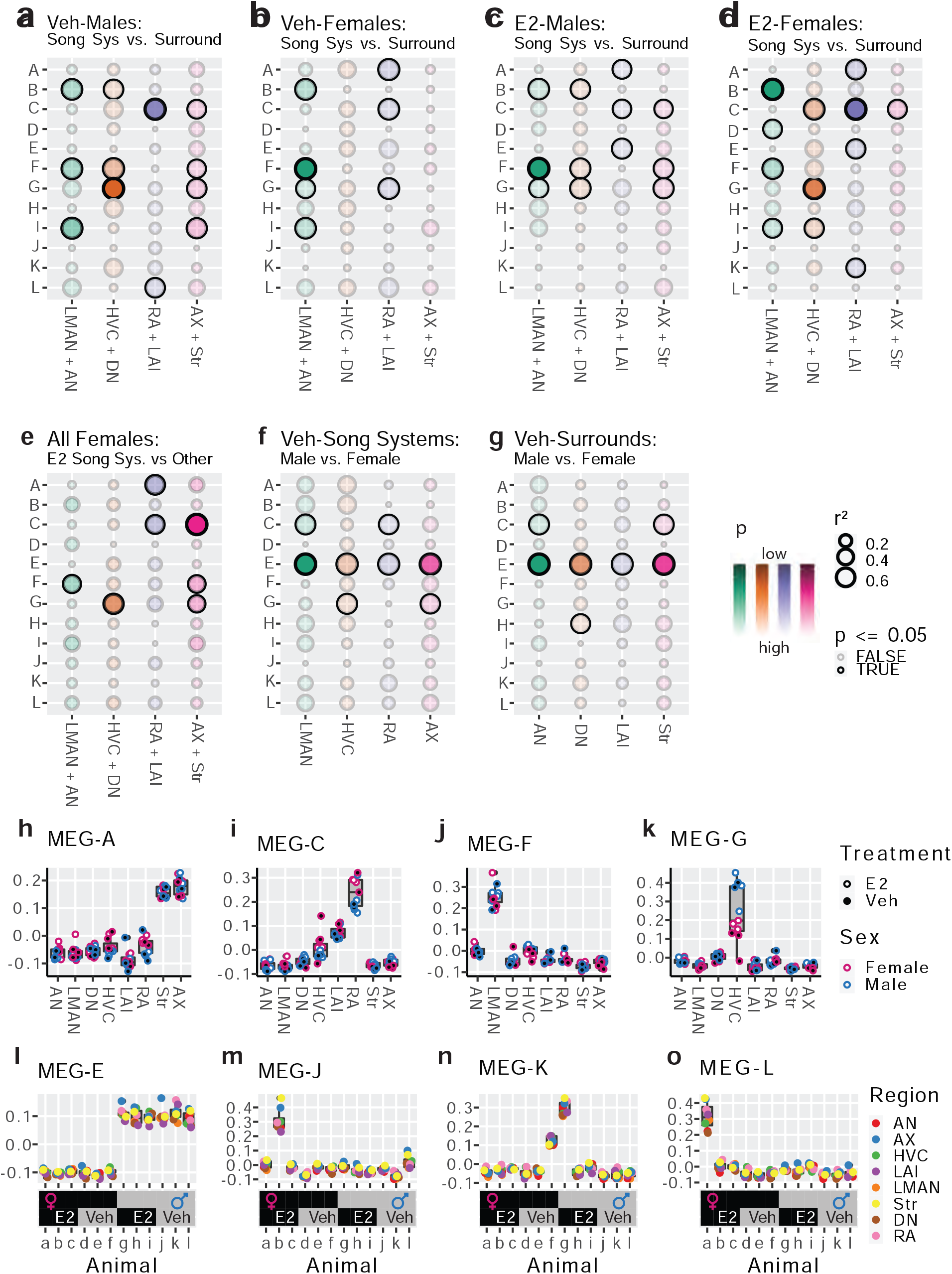
Association of modules to experimental variables. **a-g,** Bubble plots showing statistical association between MEG expression and variables of interest in various sample subsets. Strength of association (r²) is encoded by bubble size, significance (p) is encoded in the color scale with significant associations darkly bordered. Pearson correlation and Students t test, alpha=0.05. Plots show the associations between gene modules (rows) to; **a-b,** vehical treated song system specializations, comparing MEG expression in song system samples from either sex to their appropriate surrounding controls; **c-d,** E2-treated song system specialization, same comparison as **a-b** but within E2-treated samples; **e,** female vocal learning capacity after E2, comparing E2 treated female song system components to all other female samples from that circuit node; **f,** sexual dimorphism within the song system, comparing vehicle treated male and female song system components; **g,** sexual dimorphism within the surrounding control regions, comparing the vehicle treated male and female surrounding control samples. Each neural circuit node is considered separately (columns). **h-k,** Expression of modules with strong region specific expression. Module A is highly expressed in Str and Area X samples, with addtional differences between RA and LAI (**h**); Module C is highly expressed in the arcopallium, especially RA, with some increaase in HVC (**i**); Module F is expressed highly only in LMAN regardless of sex or treatement (**j**); Module G is only highly expressed in HVC, where it differs both by sex and treatment (**k**). **l-o,** Expression of selected module eigengenes by animal (top) aligned to their respective experimental variables (bottom), color indicates region. The sex chromosome enriched module E was highly expressed in all male samples and depleted in all female samples regardless of brain region or pharmacological treatment. Module J, K, and L eigengenes were each highly expressed in samples from one (J and L) or two (K) animals across all brain regions sam- pled.

### E2-responsive gene modules in song nuclei

We next assessed the effects of chronic exogenous estrogen on the developing song system. In the E2-treated juvenile males, the song nuclei specialized modules overlapped to those seen in vehicle-treated males (**Fig. 2a vs 2c**). Differences were that: in LMAN and Area X module I was no longer present; in RA module L was no longer present, module A appeared as seen in females;, and module E also appeared. These differences in gene module specializations in the male song system are consistent with the E2-treatment regimen used causing a slight decrease in vocal learning accuracy in males^29^.

In contrast, E2-treated females more closely mirrored the gene module specializations seen in the vehicle treated male song system (**Fig. 2a vs 2d**). Specifically, in LMAN, modules B, F, and I were retained similar as in vehicle-treated males, and module D uniquely appeared; in HVC, module G expression appeared and was strongly specialized as in vehicle-treated males, but modules C and I appeared relative to vehicle-treated females (Fig. 2b vs 2d); in RA, module G disappeared and module C was retained as in vehicle-treated males, and module A was retained and module K appeared relative to vehicle-treated females.; and in Area X, only module C appeared specialized in the E2-treated females. That is, the transcriptional response of female song nuclei to E2-treatment appeared to be far more dramatic than in males.

We performed an additional test for E2-induced changes to gene module expression in females by comparing E2-treated song nuclei in females to the combination of E2-treated surrounds from the same animals, vehicle-treated female surrounds, and vehicle-treated female song nuclei (**Fig 2e**). This analysis represents a comparison between samples from vocal learning capable female samples and non-vocal learning female samples at each node. It revealed 1 to 3 modules in each song nucleus of E2-treated females that were a subset of some of the strongest correlated modules found in control males (**Fig. 2a vs 2e**). These were: module F in LMAN; module G in HVC; module C in RA; and modules C, F and G in Area X. That is, these are four core modules for song nuclei that are the most sensitive to E2-induction or enhancement in females and most associated with the presence of vocal learning in females.

### Gene modules for telencephalic sex differences

We next tested whether some modules could be explained by sex differences in the brain, regardless of song nuclei presence or vocal learning status. We compared vehicle-treated male and female song nuclei and surrounds separately, and found that module E strongly correlated with higher expression in all male brain regions relative to females (**Fig. 2f,g**). Among the song nuclei, module G eigengene expression was significantly higher in male HVC and Area X; conversely, module C expression was higher in female LMAN and RA (**Fig. 2f**). Among the surrounds, module H eigengene expression (not significant in any other comparison) was significantly higher in female DN and module C was significantly higher in female AN and Str (**Fig. 2g**). These findings indicate that module E contains genes that could be explained by a broad transcriptome sex difference in the brain, whereas the expression of other modules are more specific to brain region and/or treatment.

### Gene modules with region and bird specific expression

We next quantified the magnitude of module expression differences and observed several gene modules with clear region-specific expression patterns (**Fig. 2h-k**). Module A, which appeared specialized in RA relative to LAI in several comparisons, was highly expressed in all Area X and Str samples, regardless of treatment or sex (**Fig. 2h**). Interestingly, *FOXP2*, a gene critical for spoken-language and vocal learning in humans^41^ and songbirds^42^, was a potent member of module A, correlating with the eigengene at r^2^ = 0.92 across all samples. Module C, which was specialized to RA, HVC, and Area X relative to their surrounds and sexually dimorphic in LMAN and Str, was most highly expressed in arcopallial samples, especially RA, with some increase in HVC (**Fig. 2i**). Module F, which was specialized to LMAN relative to its surround in all such comparisons, was highly expressed exclusively in LMAN regardless of sex and E2-treatment (**Fig. 2j**). Module G, which was specialized to HVC in a sex- and E2-dependent manner, was only highly expressed in HVC samples, with males having higher expression than females, and females further split by treatment (**Fig. 2k**).

We also checked if genes in specific modules were enriched in their expression in specific animals or divisions of animals regardless of region. The module E eigengene was highly expressed in all male samples and lowly expressed in all female samples regardless of brain region or treatment (**Fig. 2l**), consistent with brain sex differences (**Fig. 2f,g**). At the other extreme, three small modules showed higher expression specific to individual birds: module J was highly expressed in all samples of animal ‘b’, an E2-treated female (**Fig. 2m**); module K was highly expressed across all samples of animal ‘f’ vehicle-treated female and at roughly half-dose in animal ‘g’, an E2-treated male (**Fig. 2n**); module L was highly expressed in animal ‘a’, another E2 treated female (**Fig. 2o**). These findings indicate that the three smallest modules, J, K and L, although with some patterning to RA for the later two (**Fig. 2a,d**), are strongly animal specific in their expression. As we can think of no source of technical variation that would produce a broadly distributed shift in the neural expression of a single gene module, this variation is likely attributable to biological interindividual variation, whether this be genetic or from life history we cannot say from the present data.

### Functional enrichment of specialized modules

We sought to understand the cumulative biological function of genes among the modules. To do this, we mapped the zebra finch genes to their 1:1 human orthologs where possible and then used human gene annotation to examine the Gene Ontology (GO) functions enriched within each module’s constituent genes. Of the 12,444 module assigned genes, 7,909 (63%) had 1:1 human orthologs annotated in Ensembl. We found GO terms significantly enriched in module G (**Table 1**), which included “DNA binding transcription factor activity”, “cell differentiation”, “anatomical morphogenesis”, “cell-to-cell signaling”, and “positive regulation of multicellular organism growth”, indicating that module G genes specialized in male and E2-treated female HVC and Area X potentially act to integrate and differentiate late born neurons. Other significantly enriched terms were “extracellular matrix structural component”, “external side of the plasma membrane”, and “extracellular space”, indicating that this module may also act to restructure the extracellular matrix, perhaps to accommodate new cells. Module E, which was differential between the sexes for both song nuclei and surrounds, had 6 significantly enriched terms, of which 3 pertained to DNA damage repair: “nucleolus”, “transcription, DNA templated”, and “U2-type precatalytic spliceosome” (**Table 2**). These results in module E indicate that there is a sexually dimorphic gene expression program broadly distributed across the finch telencephalon that likely acts within the nuclear environment. A full table of GO enrichments by module can be found in supplemental data (supplemental_sig_go_enrich.csv).

**Table 1 -.** Module G functional enrichment analysis. Significantly enriched GO terms within the 1:1 human orthologs of module G. Lists full GO term, GOid, and uncorrected p value calculated by GAGE.

| GOterm | GOid | p |
| --- | --- | --- |
| DNA-binding transcription factor activity, RNA polymerase II-specific | GO:0000981 | 0.012 |
| cell differentiation | GO:0030154 | 0.013 |
| DNA-binding transcription factor activity | GO:0003700 | 0.015 |
| extracellular matrix structural constituent | GO:0005201 | 0.024 |
| inflammatory response | GO:0006954 | 0.026 |
| anatomical structure morphogenesis | GO:0009653 | 0.03 |
| external side of plasma membrane | GO:0009897 | 0.036 |
| extracellular space | GO:0005615 | 0.037 |
| DNA-binding transcription activator activity, RNA polymerase II-specific | GO:0001228 | 0.04 |
| cell-cell signaling | GO:0007267 | 0.041 |
| positive regulation of multicellular organism growth | GO:0040018 | 0.042 |
| transmembrane transporter activity | GO:0022857 | 0.043 |

**Table 2 -.** Module E functional enrichment analysis. Significantly enriched GO terms within the 1:1 human orthologs of module G. Lists full GO term, GOid, and uncorrected p value calcu-lated by GAGE.

| GOterm | GOid | p |
| --- | --- | --- |
| nucleolus | GO:0005730 | 0.024 |
| transcription, DNA-templated | GO:0006351 | 0.031 |
| DNA repair | GO:0006281 | 0.037 |
| damaged DNA binding | GO:0003684 | 0.044 |
| U2-type precatalytic spliceosome | GO:0071005 | 0.047 |
| cellular response to DNA damage stimulus | GO:0006974 | 0.048 |

### Gene modules enriched for human speech associated genes

We next asked if any of the modules were enriched for genes previously determined to be convergently specialized in songbird song nuclei and human speech brain regions^8,40^. These gene lists included the transcriptional convergence between RA and human dorsal laryngeal motor cortex (dLMC), HVC and dLMC, LMAN and dLMC, Area X and the anterior caudate, and Area X and the anterior putamen (Gedman et al. Fig3B,D^9^). We found that module B, specialized to both LMAN and HVC (**Fig. 2a**), was enriched for the convergently specialized genes in human dLMC (**Fig. 3a**) known to match the uppers layers of the cortex^9^. Module C, specialized to RA (**Fig. 2a,i**), was enriched for the genes convergently specialized in dLMC and RA (Fig 3a) known to match the lower layers of the cortex^9^. Module A, highly expressed in Area X and Str (**Fig. 2h**), was enriched for genes convergently specialized in Area X and human anterior striatum (both caudate and putamen; **Fig. 3a**). Module I, specialized to Area X (**Fig. 2a**), was also even more strongly enriched for the same convergences as module A (**Fig. 3a**). These findings indicate that the genes previously identified as convergently regulated between songbird song brain regions and human speech brain regions are components in the larger specialized gene networks identified here using WGCNA.

**Figure 3 -.**
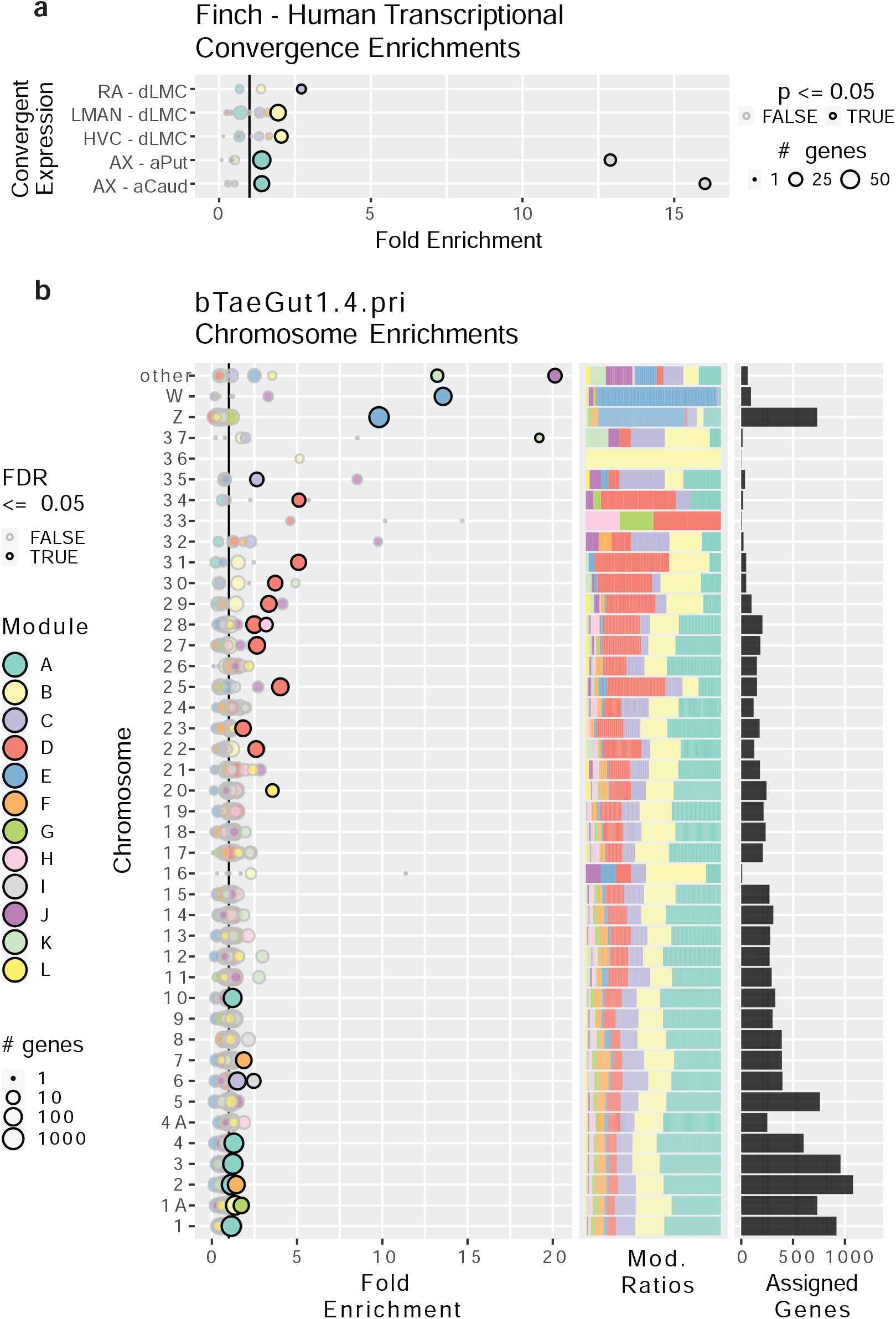
Gene module enrichments for human convergent signature and for chromosomes. **a,** Enrichment of genes previously found to be convergently differentially expressed in the human laryngeal motor cortex and the pallial song nuclei or convergently expressed between the human vocal striatum and Area X. Bubble size linearly encodes the number of genes in each convergence module pairing. Significance was assed using a one-tailed GAGE test, similar to GO ontologies, alpha=0.05. Significant enrichments are darkly bordered and opaque. Values to the right of the vertical black line indicate above random chance. **b,** Enrichment of genes from specific chromosomes. Left, fold enrichment of modules onto zebra finch chromosomes in the newest genome assembly available; center, the portion of module assigned trascripts from each chromosome per module; right, the number of module assinged genes per chromosome. Each row is a chromosome with each bubble representing the enrichment of transcripts from that chromosome in one of the gene module defined by WGNCA. Values to the right of the vertical black line indicate above random chance. The size of the bubbles indicates the log10 transformed number of genes in each chromosome module pairing. Significance was assessed using an FDR corrected boot- strapped test of observed enrichment for each module chromosome pairing based on 50,000 randomizations of genes into modules. Significant enrichments are darkly bordered and opaque. **a-b** use the same color scale for modules.

### Gene modules enriched for specific chromosomes

We next determined whether any modules were enriched in genes from specific chromosomes. We performed a bootstrapped enrichment analysis, randomizing the mapping between genes and modules 50,000 times to approximate null distributions. P values for each chromosome-module pairing were then FDR corrected. To do this as accurately as possible, we performed this analysis using revised chromosomal assignments and structure from the most recent zebra finch genome assembly, bTaeGut_1.4.pri, which better resolves the microchromosomes^43^. Surprisingly, we found that each of the 12 modules were enriched for genes from at least one chromosome (**Fig. 3b**). The most striking was module E genes, which were enrinched on the Z and W sex chromosomes, with nearly all W expressed genes and ∼⅔ of Z expressed genes being members of module E. This is consistent with module E exhibiting higher expression in male samples regardless of treatment or region (**Fig. 2d-e**). For the autosomes, we observed significant enrichments of two HVC specialized gene modules on chromosome 1A: module B (specialized in male HVC and human dLMC); and module G (specialized in male HVC and E2-treated female HVC and sexually dimorphic in vehicle-treated HVC). These results are particularly interesting given that zebra finch chromosomes 1 and 1A are believed to be the result of a songbird-specific fragmentation of the ancestral chromosome 1 found in chickens^44,45^.

Module A, the largest module, which was highly expressed in Area X and adjacent striatum, was enriched across 5 macro-chromosomes (chr1, 2, 3, 4, and 10). Module F, which was strongly expressed in LMAN, was enriched on chromosomes 2 and 7. Modules C (most strongly expressed in the arcopallium and part of RA and Area X specializations) and I (a component of the LMAN specialization) were enriched on chromosome 6 and microchromosome 35. Module D, a component of the LMAN specialization in E2-treated females, was enriched across 8 small microchromosomes (chr22, 23, 25, 27, 28, 29, 30, 31). Most (4 of the 5) of the smallest modules in gene counts were enriched in the microchromosomes: Module H, which was sexually dimorphic only in DN, was enriched on microchromosome 28; module K was significantly enriched on microchromosome 37 and in the “other” category, which includes all remaining unnamed DNA scaffolds in the assembly such as further microchromosomes; module J was also enriched in “other”.

### Sex chromosome gene expression across regions

To better understand the relationship between the sex chromosomes and module E, we examined the distribution of membership in module E with the sex chromosomes separated. WGCNA allows us to consider gene membership in a module as a continuous variable, rather than a binary variable, by correlating each gene’s expression profile to the module eigengene. Doing this for module E, we found that Z transcripts were positively correlated to the module eigengene while W transcripts were anticorrelated (**Fig. 4e**). This is consistent with Z chromosome transcripts were generally lower expressed in females relative to males, while W chromosome transcripts were only expressed in female brains. This general reduction in female Z chromosome transcript abundance within module E is consistent with the finding that diploid Z chromosomes in male birds do not have one copy inactivated to compensate for gene dosage, unlike the X inactivation in female mammals^34,46^.

**Figure 4 -.**
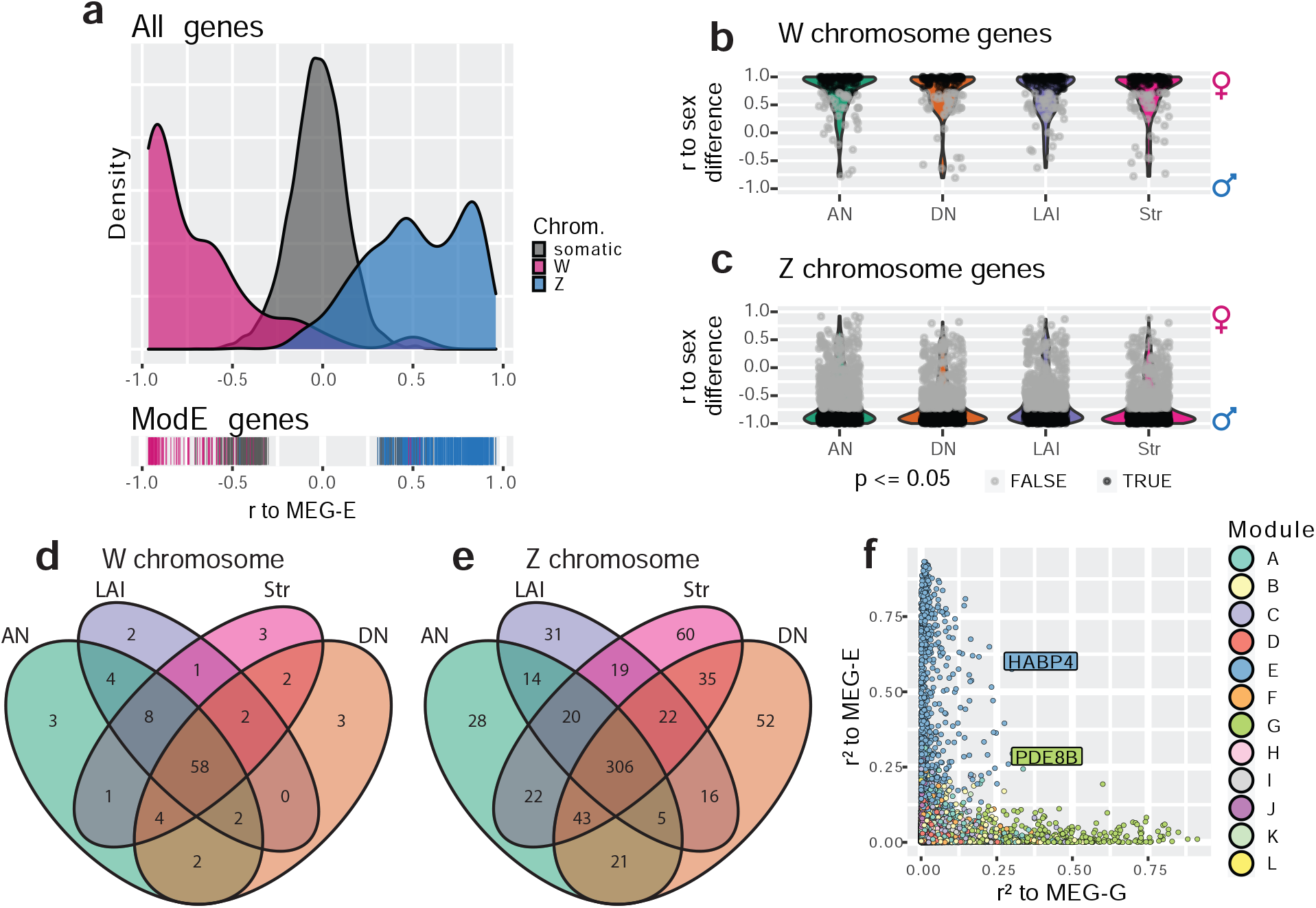
Brainwide signatures of sex chromosome expression. **a,** Distribution of continuous membership in module E across all module assigned genes (top) and module E assigned genes (bottom) based on corelation of expression to the module eigengene (Pearson r to MEG-E) with sex chromosomes separated. **b-c**, Distribution of sex chromosome gene expression correlations to the sex difference in vehicle treated finches. Postive correlations indicate female biased expression while anticorrelations indicate male biased expression. Significance was assessed in each region using an upper-tailed student’s correlation test for W chromosome transcripts (**b**) and lower-tailed for Z chromosome transcripts (**c**), with significant correlations in black, alpha=0.05. **d-e,** Venn diagrams intersecting the significantly sex difference correlated genes across non-vocal surround regions for the W and Z chromosomes respectively. **f,** Comparison of continuous membership in module E (r2 to MEG-E, y-axis) and module G (r2 to MEG-G, x-axis) across all 12,444 module assigned genes.

Given the sexually dimorphic expression of module E across brain regions and the enrichment of the sex chromosomes within that module, we separately compared the expression of sex chromosome genes in the non-vocal motor surrounds of vehicle-treated control males and females regardless of WGCNA assignment to better understand sex chromosome expression without the influence of vocal learning specialization or E2 response. To do this, we first tested for correlated expression between each sex chromosome transcript to the animals’ sex within each region such that female-enriched transcripts were positively correlated and male-enriched transcripts were anticorrelated with sex (**Fig. 4f,g**). In these gene sets, we identified between 73 and 82 significantly expressed W chromosome genes and between 433 and 527 significantly depleted Z chromosome genes for each of the surround regions in males relative females. Examining the union of these gene sets revealed that a total of 95 W and 694 Z chromosome genes were differentially expressed between sexes in at least one brain region, 62% and 65% of annotated sex chromosome genes respectively. Conversely, the intersection of these regional gene sets contained 58 significantly expressed W chromosome genes and 306 significantly depleted Z chromosome genes in all non-vocal regions in females, representing 38% and 29% of annotated sex chromosome genes, respectively (**Fig. 4h-i**, **Table S2**). This indicates that there is both a large broadly distributed set of sex enriched/depleted sex chromosome genes and regional patterned sex chromosome gene expression.

### Modeling vocal learning and sex chromosome module interactions

As it is unlikely that the gene modules act independently of each other, we sought putative interacting genes between two modules of interest, module G specialized in HVC and Area X and module E dominated by sex chromosome genes. To do this, we again replaced binary in-or-out module membership with continuous module membership by correlating each genes’ expression to the relevant module eigengene^38^. This allowed us to quantify the extent to which any gene was associated with any module, regardless of initial assignment^33^. Looking across all assigned genes for our modules of interest, we identified two outlier genes, *PDE8B* and *HABP4*, which were the most E associated gene assigned to module G and the most G-associated gene assigned to module E, respectively (**Fig. 4j**). Both genes were significantly upregulated in male HVC relative to its surround, but not in female HVC of either treatment. *PDE8B* catalyzes the hydrolysis of the second messenger cAMP and mutations to the gene cause an autosomal dominant form of striatal degeneration in humans^47^. *HABP4* is an RNA binding protein, known to repress the expression and subsequent DNA binding of *MEF2C*^48^, a Z chromosome transcription factor which has undergone accelerated evolution in songbirds^49^ and whose repression by *FOXP2* is critical for cortico-striatal circuit formation in mice related to vocal behaviors^50^. Both *PDE8B* and *HABP4* are found on the Z chromosome and *HABP4* was one of the 694 Z transcripts significantly reduced across all brain regions in vehicle-treated females relative to males (**Table S2**). These findings again indicate that Z chromosome genes may be subject to multiple gene regulatory programs: the broadly distributed brain transcript reduction driven by reduced sex chromosome copy number; and the specialized upregulation in HVC for vocal learning behavior.

### Sex chromosome dosage effects by module

To better understand the relationship between gene modules, sex chromosome gene expression levels, sex chromosome dosage, and song nuclei specializations, we directly compared the abundance of Z and W transcripts between vehicle-treated male and female samples in surrounds and song nuclei. After averaging across samples for each brain region, and combining them for all brain regions, Z chromosome transcripts were present in both females (pink) and males (blue) as expected (**Fig. 5a**). Although more Z chromosome genes were assigned to module E, many were assigned to the other modules (**Fig. 5a**), consistent with our chromosome mapping without enrichment analysis (**Fig. 3b**). However, there were clear male vs female expression differences apparent for Z chromosome genes in module E, but no obvious sex differences for Z chromosome gene expression in other modules. In contrast, W chromosome transcripts were mainly expressed in females (pink) and not males (blue), and were generally restricted to module E (**Fig. 5b**), also consistent with our chromosome enrichment analysis (**Fig. 3b**). To further quantify these effects outside and inside of the song system, we computed the percent of total expression for each sex chromosome gene which came from male samples (male_avg/(male_avg+female_avg)) and compared these distributions to those predicted by chromosome dosage. The expected average percentages based upon dosage are 66.6% male expression for Z chromosome genes (2 Zs in males vs. 1 Z in females; **Fig. 5c-f**, red lines) and 0% male expression for W chromosome genes (0 W in males vs 1 W in females; **Fig. 5g-j**, red lines).

**Figure 5 -.**
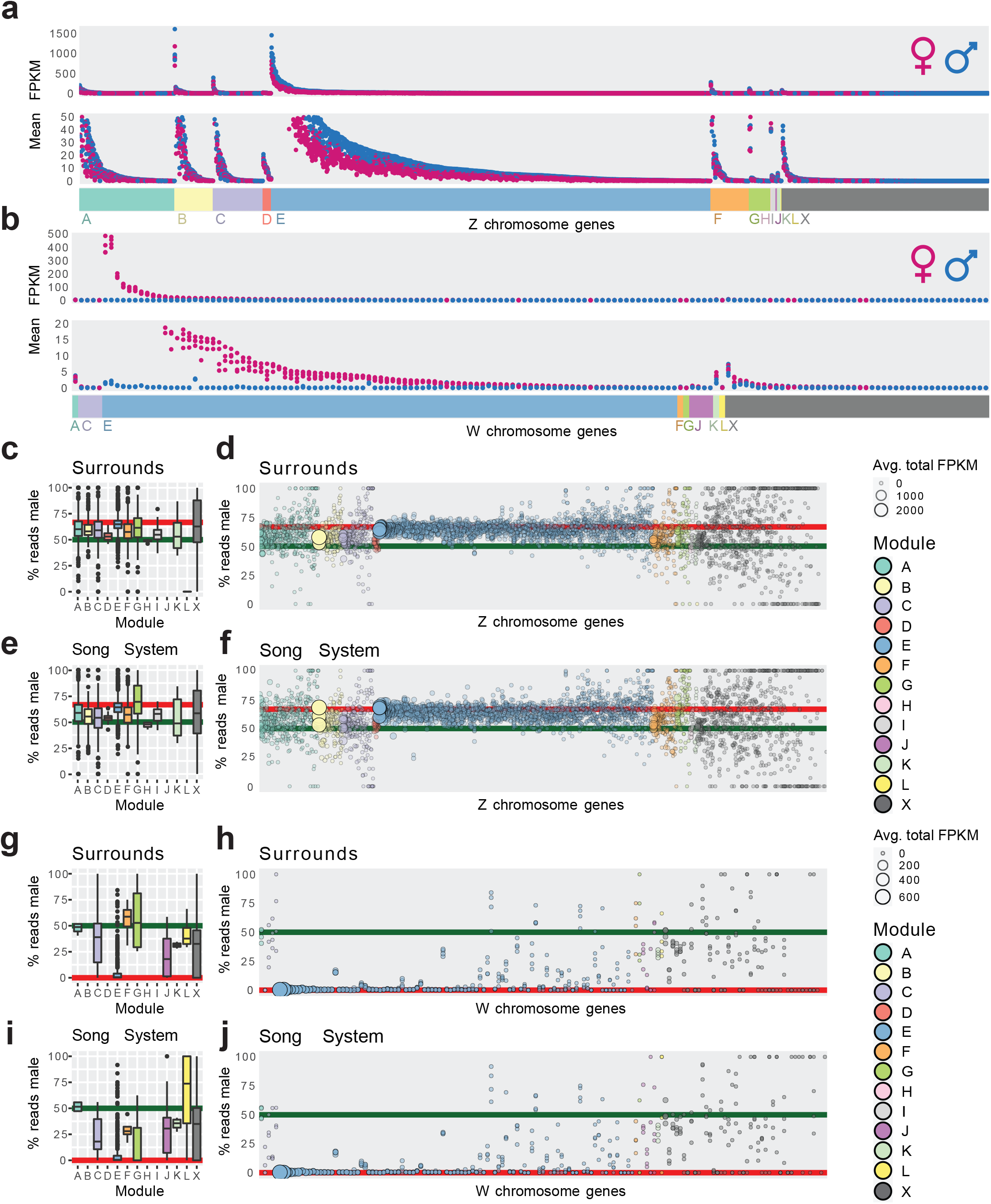
Sex chromosome dosage effects by module. **a,** Scatter plot of transcript expression abundance for all Z chromosome genes (y-axis) ordered by module assignment and then expression (x-axis). Each dot is one gene with expression level measured as the average of one brain region in vehicle-treated male birds (blue, n= 3)) or female birds (pink, n = 3)). The upper graph shows all genes while the lower graph has a cut-off y axis to better show lowly expressed genes. **b,** similar as **a**, except for all W chromosome genes. **c,** Boxplots showing the distribution of the percent reads from male surrounds per Z chromosome gene from **a** (y-axis) and their module assignment (x-axis). Red line indicates the male read percentage expected for Z chromosome genes, 66.6%; green line indicates equal expression between sexes. **d,** Bubble plot of the underlying Z chromosome single gene data from **c** X-axis is ordered according by module assignment, encoded by bubble color, and expression. Bubble size and opacity indicates cumulative average expression (male avg. FPKM + female avg. FPKM), with higher expressed genes being larger and more opaque. **e-f,** same as **c-d** but for song nuclei. Note that the Z chromosome genes of module E are expressed on average at almost exactly the sex ratio predicted by dosage while Z chromosome genes in other modules show some degree of compensation. **g-j,** same as **c-f**, except for W chromosome genes. Red line indicates the male read percentage expected for W chromosome genes, 0%; green line indicates equal expression between sexes.

Module E assigned Z chromosome genes were expressed at almost exactly the ratio predicted by Z chromosome dosage on average; 65.1% measured versus 66.6% predicted, and the most abundantly expressed transcripts (largest circles) were expressed closest to the dose prediction in both surrounds (**Fig. 5c,d**) and song nuclei (**Fig. 5e,f**). For non-E modules, their Z chromosome transcripts were intermediate between equal expression and the prediction from chromosomal dosage in surrounds (**Fig. 5c,d**), with Z chromosome genes in module D and H were the least male biased across regions (**Fig. 5e,f**). One major difference between surrounds and song nuclei was the Z chromosome gene expression levels again in module G. The 25 Z chromosome genes in module G were more male biased in song nuclei compared to surrounds (**Fig. 5c,d vs e,f**) and were the only set of Z chromosome genes that were male biased above the predictions from dose on average (**Fig. 5e**). These findings indicate that the expression levels of Z chromosome genes in module E were predominantly dose regulated, with the abundance of the RNA dictated by the presence versus absence of those specific chromosomes regardless of brain region sampled. However, Z chromosome genes in the other modules whose expression is enriched for one or several brain regions, animals, or treatments exhibited varying degrees of dosage compensation to overcome the Z chromosome difference.

In contrast, W chromosome genes assigned to module E had on average 6.8% male read mapping counts (median 0.5%; **Fig. 5g,h**). Non-module E assigned W genes had 33.0% male reads mapped (median 32.7%; **Fig. 5g,h**). As the values should be 0% reads from males mapped to the W chromosome, we believe this male read mapping is from less divergent transcript paralogs from the Z chromosome that map to the W chromosome. Examining the data at the level of single genes, we observed that module E assigned W genes were far more likely than non-module E genes to show no- or little- putative Z paralog mapping (more genes on or near the 0% line; **Fig. 5g,h**). Within module E, the genes with the lowest expression (smaller circle size) were the most likely to have putative paralogous expression (the lowest ∼⅓ of these genes contributed to the majority of expression above 0%, **Fig. 5h**). We observed similar results in module E versus other modules for W expression in song nuclei (**Fig. 5i,j**). While the male read mapping of the W genes in modules F, G, and L do appear to shift in the song system relative to the surrounds, each of these modules contains a single W gene and all three of these genes are lowly expressed (**Fig. 5b**).

These results demonstrate that rather than being confounded by chromosome dose, WGCNA allowed us to resolve the effects of dose in an unbiased way. Module E grouped together the un-dosage-compensated Z chromosome and W chromosome genes across brain regions. In contrast, the Z chromosome genes placed in modules specialized for one or more song nuclei had some level of dosage compensation in males. This compensation appeared regionally patterned for Z chromosome genes in module G (enriched in 2 or more song nuclei depending on treatment), which were specialized beyond the normal chromosome dosage only within the song system. Average expression values of all sex chromosome genes for each brain region from vehicle-treated animals can be found in supplemental_z_chromosome_express.csv and supplemental_w_chromosome_express.csv.

### Candidate gene drivers of HVC specialization in E2-treated females

To reduce the 344 genes in module G to the putative drivers of HVC development, we next examined the relationship of membership in module G (correlation to MEG-G) to gene expression specialization in HVC at the level of single genes. We did this by testing for correlations between individual gene expression and the specialization of HVC in males (vehicle- and E2-treated) or E2-treated females in each of the following four comparisons: 1) male HVC specialization relative to the surround; 2) E2-treated female HVC specialization relative to the surround; 3) male HVC relative to female HVC in vehicle-treated controls; and 4) E2-treated female HVC specialization relative to vehicle-treated female HVC. For all four comparisons, we found the higher the correlation of module G genes to the MEG-G, the higher the correlation with the vocal learning specialization (**Fig. 6a-d**). This means that their expression was higher in male or E2-treated female HVC relative to the appropriate non-vocal learning controls across all comparisons. We identified genes of interest as being strongly correlated (r^2^ ≥ 0.5) to both the module G eigengene and vocal learning specialization (**Fig. 6e-h**, higher magnification view of colored boxes in **Fig. 6a-d**). All genes of interest exhibited a positive correlation with song nuclei gene expression specializations across all four comparisons (**Fig. 6a-d**). We next generated a core gene list from module G that correlates with both in E2- and sex-dependent HVC expression by intersecting these four vocal learning specialized gene sets (**Fig. 6i**). We found a core set of 14 genes that strongly marked vocal learning capable HVC in all comparisons and strongly correlated with the aggregate of module G expression (**Table 3**, **Fig. S6**). The results of each individual comparison can be found in supplemental_hvc_specializations.xslx.

**Figure 6 -.**
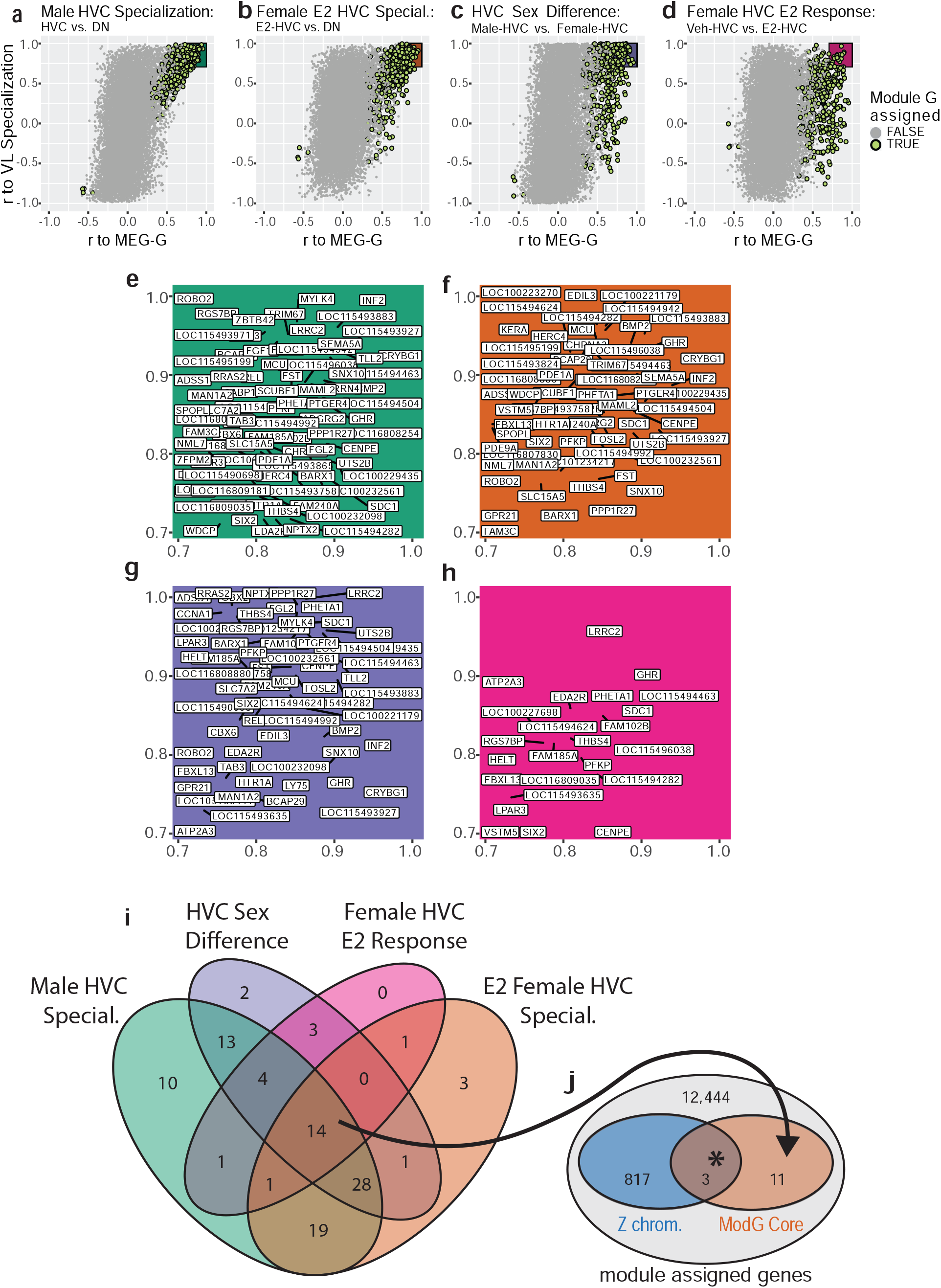
Identification of core genes in module G and their association to the Z chromosome. **a-d,** Single gene continuous membership in module G (x-axis; Pearson r to MEG from module G) for all assigned genes vs correlation to vocal learning in masculine or mas-culinized HVC relative to samples from non-vocal learning females in each of the four com-parisons; **a,** male song system membership, comparing individual gene expression in male HVC samples of either treatment to expression in the surrounding DN; **b,** female vocal learn-ing capacity after E2, comparing E2 treated HVC to all other female DN or HVC samples; **c,** sexual dimorphic gene expression within the song system, comparing vehicle treated male and female song system components; **d,** estradiol responsive gene expression in female HVC, comparing E2 treat and vehicle treat female HVC samples. Each point is a gene colored by module assignment, the shaded area indicates gene of interest criteria for each comparison. **e-h,** Blowup of shaded regions in **a-d** respectively showing genes of interest from each comparison. **i,** Identification of core genes by intersecting the four gene sets of interest. **j,** Enrichment of Z chromosome transcripts within the core genes. * indicates p = 0.0087 by an upper- tailed hypergeometric test.

**Table 3 -.** Core genes of module G specialization to vocal learning HVC. Putative drivers of module G specialization to vocal learning capale HVC. Intersection of the four gene of interest sets (Fig. 4b), separating the significant enrichment of genes from the Z sex chromosome.

| Z Chromosome | Other | cont. |
| --- | --- | --- |
| GHR | CENPE | LOC115494624 |
| LRRC2 | EDA2R | PFKP |
| RGS7BP | FAM102B | PHETA1 |
| THBS4 | FBXL13 | SDC1 |
|  | LOC115494282 | SIX2 |
|  | LOC115494463 |  |

We examined the chromosomes of origin for these 14 core genes and found that three (*GHR*, *RGS7BP*, and *THBS4*) were on the Z sex chromosome (**Fig. 6j**). This was a >3-fold significant enrichment over chance of Z chromosome transcripts (p = 0.009, upper-tailed hypergeometric test) against a background of module assigned genes. This result was statistically significant regardless of the background gene set, when using all genes (∼21,000) or only module G members (344). These three Z chromosome module G genes in HVC not only exhibited >50% reduced expression in control female HVC relative to males, but also exhibited upregulated expression in male HVC relative to the surrounding DN regardless of treatment and upregulation in E2-treated female HVC relative to either the surround or vehicle-treated female HVC (**Fig. S6**). In the context of our cross-region sex chromosome analysis (**Fig. 3c-f**), only *RGS7BP* was sexually dimorphic outside of the song system, being absent in all female surround regions. Neither *GHR* (growth hormone receptor) nor *THBS4* were significantly depleted in any surround region comparison. Taken together, these results indicate that the 3 core genes in vocal learning capable HVC on the Z chromosome are subject to additional E2 sensitive transcriptional regulation in HVC, separate from the Z chromosome transcript reduction seen throughout the female brain.

Of the 14 core genes total, several have been previously studied in the brains of other species which may inform their role in vocal learning systems. *THBS4* encodes a secreted extracellular-matrix glycoprotein necessary for appropriate neuronal migration in the mouse^51^ and is elevated 6-fold in the human cortex compared with non-vocal learning primates^52^. Human *EDA2R* was recently identified as a top correlate of cognitive performance and brain size^53^; it was also found in a human GWAS study that correlated it with circulating estrogen and testosterone levels^54^. Rare mutations in human *PHETA1* lead to Lowe oculocerebrorenal syndrome, that includes pathophysiology in seizures, mental retardation, and structural brain abnormalities^55,56^. *SIX2* is a homeobox domain containing transcription factor that governs early brain and craniofacial development and provides neuroprotection from dopamine injury^57,58^. *GHR* encodes a transmembrane receptor whose activation controls cell division^59^. The gene which encodes GHR’s ligand, growth hormone (GH), is interestingly duplicated and undergoing accelerated evolution in the genomes of songbirds, is upregulated in the zebra finch auditory forebrain following the presentation of familiar song, and exerts anti-atrophic influence during chicken development^60–62^. Based on these findings, we consider *GHR* as the most likely candidate gene related to the E2 sensitive atrophy of HVC in females.

## Discussion

The present study seeks to further our understanding of sexually dimorphic vocal learning in zebra finches by comparing the gene expression specializations in the song system between male and female birds at PHD30 and in response to E2-treatment, at the onset of atrophy of the female vocal circuit. The birds were given either a vehicle or E2 from hatching, which induces rudimentary vocal learning behavior in females where it would otherwise be absent. In control females, HVC appeared unspecialized at the level of gene module expression, with no significantly differentially expressed MEGs compared to the surrounding nidopallium. However, in E2-treated females, HVC exhibited a subset of the observed male HVC gene expression specializations. Similarly, in the vehicle-treated females, the striatum located where Area X would be also lacked any specialized gene module expression, but the E2-treated female Area X had a subset of specialized gene expression as in males. This contrasts with RA and LMAN which were similarly specialized in males and females in the absence of E2-treatment. Given that lesions of HVC prevent the emergence of Area X in E2-treated females^32^, these results support a model of zebra finch development where transcriptomic masculinization of female HVC by E2 is a critical event which facilitates the emergence of female Area X and ultimately endows these females to produce some rudimentary learned song.

How did E2 treatment produce the transcriptomic effects we observed with module G in female HVC and what are the implications regarding sexually dimorphic vocal learning in the zebra finch? One possibility is that this process begins with increased estrogen receptor (ER) activity within developing HVC cells after being provided surplus activating ligand^63^, followed by altered transcription of ER targets in the genome. Module G contained the androgen receptor (**Table S1**), which is believed to be a major downstream effector of the E2 response in female zebra finches^33^. This initial transcriptional loading of the system would then have been processed by gene regulatory networks within each cell, spreading in effect through the transcriptome. It is possible that differential module G expression arose purely from traditional gene regulatory networks, where transcription factors form complex, elaborate feedback networks with themselves and the genes they regulate. However, this framework fails to explain why the core 14 vocal learning correlated genes from module G in HVC were enriched for transcripts from a single chromosome: the Z sex chromosome with halved copy number in females.

We hypothesize that the Z chromosome genes identified here are co-regulated, necessary components of a growth-enabling transcriptional program downstream of ER, represented by module G. This module G transcriptional program could be specialized to developing male HVC by the expression of patterning genes, such as *SIX2,* early in development and maintained through persistent *GHR* signaling. We propose that these Z chromosome transcripts in module G are reduced in females by lower haplotype dosage during development and thus fail to specialize female HVC. Due to lower abundance of gene products from module G, female HVC may be unable to accommodate new neurons during juvenile development and fail to facilitate emergence of its downstream target Area X. Similarly, without a sufficiently developed HVC, RA lacks one of its major inputs and may subsequently atrophy. We propose that E2 masculinizes female song behavior by increasing the abundance of these module G transcripts in HVC, increasing specialized HVC growth, and facilitating emergence of HVC’s other major target, Area X (**Fig 7**). This model of sex chromosome influenced song system development is consistent with recent work comparing male and female zebra finch transcriptomes from RA at young juvenile (PHD20) and young adult (PHD50) ages in un-manipulated birds (Friedrich et al. 2022)^64^. While that study proposed that the role of the sex chromosome in maintaining transcriptomic sex differences diminishes across development as the proportion of specialized genes that originate on the sex chromosomes diminishes, this effect was driven by large increases in differentially expressed autosomal genes rather than by any reduction in sex chromosome dimorphism; the percentage of differentially expressed Z chromosome genes increased from 28% at PHD20 to 39% at PHD50^64^. This leads us to conclude that sexually dimorphic Z chromosome expression in juveniles precedes the sexually dimorphic expression of the autosomes seen in adults. This is consistent with our hypothesis that sufficient expression of select Z chromosome gene products (GHR, etc..) is necessary for subsequent autosomal song system specializations (module G). Further, our results are consistent with GH’s known role in avian neuroprotection, with elevated signaling associated with the survival of chicken neurons during rounds of pruning in the developing retina^65^.

**Figure 7 -.**
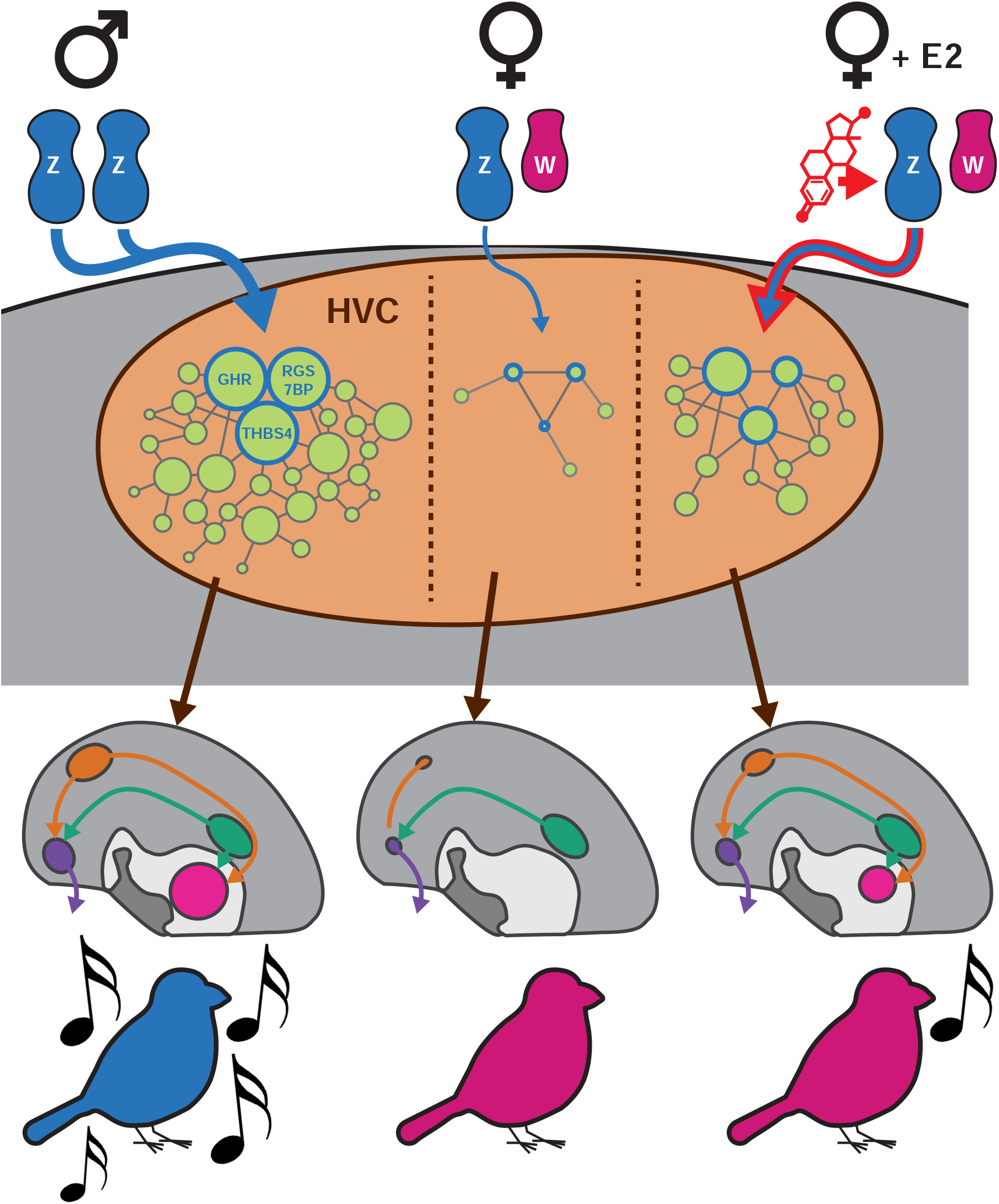
Proposed model of sexually dimorphic zebra finch vocal learning. We propose that estradiol treatment in female zebra finches masculinizes song behavior by overcoming insufficient Z sex chromosome dosage in HVC to increase the expressed of transcripts normally depleted in females. The Z chromosome genes upregulated by E2 are components in a larger proliferative genetic program which prevents HVC atrophy in males and allows for its expansion late in development. The upregulation of these genes allows for the increased specilization of the gene networks they participate in, promoting HVC development sufficiently to enable rudimentary vocal learning in females.

Our results help refine the traditional notion that hormonal signaling organizes the brain to produce sexually dimorphic behaviors independent of neuronal sex chromosome content^66,67^. Instead, our data indicate that sexual dimorphism of the zebra finch song learning ability was likely established by the interaction of sex hormone signaling and sex chromosome gene expression within HVC during development. These findings are thematically similar to work in the Four Core Genotypes mouse model^68^, where chromosomal and gonadal sex are separable by translocating the sex determining *Sry* gene. In these mice, sex chromosome composition regulates sexually dimorphic brain gene expression, circuit anatomy, and behaviors^68–71^, though the sex chromosome genes responsible remain unknown. Additional experiments manipulating the candidate genes implicated here in developing HVC to both mimic and prevent the action of E2 in female zebra finches are needed to test these hypotheses.

In addition to these vocal learning focused results, performing unbiased gene network analysis with samples taken throughout the zebra finch telencephalon in both sexes revealed a surprisingly strong relationship between the modular organization of telencephalic gene expression and the chromosomal structure of the genome. Each of the 12 modules identified by WGCNA were enriched for genes from at least one chromosome. Of these significant enrichments, those on the micro- and sex- chromosomes stood out as particularly strong. Module E encompassed the majority of W and Z chromosome genes and was sexually dimorphic in its expression in all sampled regions. Indeed, we found that roughly ⅓ of sex chromosome transcripts are significantly sexually dimorphic in all non-vocal brain regions, and roughly ⅔ are sexually dimorphic in at least one region. This half-brain-wide and half-regionally-patterned sexual dimorphism observed from the sex chromosomes may act as a substrate for the evolution of sexually dimorphic behaviors generally, with brain-wide shifts in expression providing a transcriptional background for evolution of sex-specific patterning in additional sex chromosome genes. These regionally patterned sex chromosome genes could then recruit gene expression networks from somatic chromosomes, resulting in the sex specific anatomical patterning of gene expression from throughout the genome. This general model is consistent with local reduction of GHR and other Z chromosome transcripts in the developing HVC lineage, leading to a loss of specialized module G expression in female HVC from somatic chromosomes. This hypothesis of an interaction between sex chromosomes and autosomes linked to the gain and loss of vocal learning in one sex can be tested in future studies.

## Supporting information

supplemental_sig_go_enrich.csv

supplemetal_w_chrom_express.csv

supplemetal_z_chrom_express.csv

supplemental_gene_module_assignment.csv

supplemental_hvc_specializations

## Acknowledgments

Funding for this work was provided by the Howard Hughes Medical Institute (EDJ), NIH-NIDCD R01-DC016224 (HM), and the NSF-GRFP (MHD). We thank Gregory Gedman, Giulio Formenti, Caitlin Gilbert, Lindsey Cantin, César Vargas, Chul Lee, and Jason Manley for conversations during visualization and analysis. We also thank Alipasha Vaziri and Tobias Nöbauer for providing the computing infrastructure used throughout. This analysis is a memorial to Mark Konishi (1933-2020) whose work on this topic influenced us greatly.

## Author contributions

MHD performed all analyses, generated all plots, and wrote the paper; HNC gathered all data and edited the paper; HM and EDJ edited the paper.

## Materials & Correspondence

Correspondence to Matthew H. Davenport and Erich D. Jarvis

## Materials and Methods

### Animal Handling and Sample Preparation

The 96 samples used in the present analysis are the E2 or vehicle treated subset of a previously published RNAseq dataset^29^. We briefly re-describe our methodology here. All animal procedures were approved by the IACUC of Duke University.

E2 (Sigma E1024-1G) was dissolved in DMSO (100mg/mL) and then diluted in olive oil (1mg/mL). 30-50uL of E2 sample or DMSO only vehicle was applied to the flank of male and female zebra finches daily from PHD0-PHD14 and on alternating days from PHD15-30 (n=3 per sex-treatment combination). We have previously shown that this treatment program is sufficient to induce song system masculinization in E2 treated female zebra finches^29^.

On PHD30, animals were sacrificed following one hour of dark isolation. Animals were anesthetized by isoflurane inhalation and rapidly decapitated. Brain hemispheres were dissected, embedded in OCT, and flash frozen in an ethanol and dry ice slurry. Sections were taken from the right hemisphere coronally at 14um onto polyethylene naphthalate (PEN) membrane slides for RNA isolation and adjacent sections taken on charged glass slides for histology or in-situ hybridizations. From the PEN membrane slides, song nuclei and surrounding control regions were laser capture microdissected (LCM) using an ArcturusXT LCM system (Nikon) guided by a Nissl stained tissue series for each animal. No sample pooling was performed, each sample originated from a single bird. This is 8 samples worth of RNAseq data per bird for 12 birds, providing 3 samples per sex-treatment-region combination, roughly a terabyte of read data.

RNA was extracted from the LCM isolated tissue samples using the Arcturus Picopure kit (Applied Biosystems KIT0204) following manufacturer’s instructions. RNA quality was assessed using an Agilent 2100 bio-analyzer and the RNA 6000 pico kit (Agilent 5067-1513). Next, cDNA was synthesized using the SMART-Seq v4 Ultra Low input RNA Kit (Takara 634892). Sequencing libraries were made with the NEBNext Ultra II DNA Library Prep kit (New England Biolabs E7645L) and cleaned-up using SPRIselect beads (Beck-man Coulter B23317). Libraries were sequenced by Novogene Co., Ltd. on the Novaseq 6000 platform (Illumina) and S4 flow cells resulting in 150bp paired-end reads.

### RNAseq read mapping and quality control

RNAseq reads were first trimmed to remove adapters and low quality base calls using Trimmomatic^72^ and then mapped to a high-quality Vertebrate Genomes Project (VGP) female zebra finch nuclear genome (bTaeGut2.pat.W.v2, GCF_008822105.2)^37^ using STAR (v2.7.1)^73^. Uniquely mapped reads were then tallied at the level of genes using Rsubread::featureCounts (R-3.6.3)and then counts normalized to fragments per kilobase of transcript per million mapped reads^74^. Multi-mapped reads were rejected as they have a higher probability of being associated with technical artifacts of sequencing or genome assembly. Read based quality control was performed with FastQC (Babraham Bioinformatics) with reports prepared by MultiQC (Python-3.5.5)^75^. This workflow was automated by the CountMatrix pipeline (https://github.com/mattisabrat/CountMatrix/). We next removed two outlier samples (one male vehicle HVC and one female vehicle RA) based upon hierarchical clustering of the sample space before computing gene-to-gene correlations (**Fig. S1**).

### Gene Module Identification

All remaining analyses were completed in R-4.2 unless otherwise specified. Data was wrangled in the tidyverse, and custom visualizations produced with ggplot, ggdendro, VennDiagram, RColorBrewer, and ggpubr^76,77^. Unsigned topological overlaps between genes (gene-to-gene correlations) were calculated in a single block with WGNCA::blockwiseModules with a soft thresholding power (β) of 6 based on scale free topology fit and model connectedness as described in the WGNCA vignette (**Fig. S2)**^38^. Next we determined an appropriate WGCNA parameterization quantitatively and qualitatively by sweeping the module size and tree cut height parameters of WGNCA::recutBlockwiseTrees. We selected our model for analysis by examining the resulting gene assignments and sample-sample distance matrices. We were able to increase or decrease the proportion of genes assigned to WGCNA modules by lowering or raising the minimum module size parameter, respectively. We found that setting the minimum size parameter below 100 genes included more genes, but with models that increasingly over-fit single samples, producing obvious outliers in the distance matrix (**Fig. S3**). Correspondingly, raising the minimum size parameter beyond 100 included fewer genes, but did not greatly reduce the number of outlier samples. Based on this we selected 100 as the minimum module size, parameterizing to explain as much transcriptomic variance as possible while minimizing the number of technically overfit samples (**Fig. S2**-**4**).

### Module association to vocal learning

Module eigengenes (MEGs) from each module were correlated against binarized song system membership, vocal learning capability, or sex and the statistical significance of each correlation assessed using WGCNA::corPvalueStudent. This was done in the following sample subsets by node: male samples broken out by treatment; female samples broken out by treatment; all female samples; song system components from either sex treated with vehicle; and surrounding control regions from either sex treated with vehicle. Within each of the 4 sex-treatment combinations we compared the song system components to surrounds at each node. Within all female samples we compared the vocal learning capable E2-treated song system elements to all other female samples from each node. Within the vehicle song systems and vehicle surrounds we compared between male and female finches for each region.

### Module gene ontology and convergent vocal learning gene expression signature enrichment

Module assigned zebra finch genes were mapped to their 1:1 human orthologs where possible, dropping unmapped or multi-mapped genes, using orthofindR::getOrthos (https://github.com/ggedman/orthofindR) which wraps Ensembl’s biomaRt. Uncorrected p values for the enrichment of human gene ontology terms within the human orthologs of module G were calculated using generally applicable gene set enrichment (GAGE) implemented in gage::gage^78^. To determine if the genes previously shown as convergently differentially expressed in the zebra finch song system and human speech brain regions mapped to specific modules, we treated these five gene lists identically to GO terms and tested for their significant enrichment across the human orthologs of each module also using GAGE.

### Analysis of sex chromosome gene expression independent of vocal learning or E2 treatment

Sex chromosome transcripts, regardless of WGCNA module assignment, were examined in the vehicle-treated non-vocal learning producing surround samples for each node. To find the most consistently expressed and depleted W and Z chromosome genes respectively, we correlated expression of each sex chromosome transcript with sexual dimorphism within each region, such that expressed W genes would be positively correlated and depleted Z chromosome genes would be anticorrelated. We computed correlations and p values using the WGCNA corAndP function; upper-tailed for the W chromosome and lower-tailed for the Z chromosome. Genes significantly expressed or depleted across regions were then intersected to identify consistently regulated transcripts across the telencephalon.

### Module enrichment on chromosomes

To associate modules to chromosomes, we bootstrapped FDR corrected p values for the enrichment of each chromosome-module pairing by randomizing the mapping of genes to modules 50k times and calculating the fold-enrichments observed on each chromosome from each module in each randomization to empirically determine the null distributions. The calculation of bootstraps and p values was performed in Python-3.5.5 and parallelized using joblib’s Parallel.

### Identification of core module G genes in HVC and Z chromosome enrichment

We defined genes of interest as the subset of significantly (T test for correlation: p≤0.05) vocal learning capability correlated genes in HVC whose expression correlated to module eigengene G (MEG-G) across the dataset at r^2^ ≥ 0.5 and to vocal learning at r^2^ ≥ 0.5 in at least one of the four vocal learning comparisons in HVC, calculated using WGCNA::corAndP. These comparisons were: all male HVC samples against all male DN samples; female E2 treated HVC against all other female samples at the node, including vehicle treated HVC; Vehicle treated male HVC against vehicle treated female HVC; and E2 treated female HVC against vehicle treated female HVC. We defined core genes as those meeting this criteria for all four HVC vocal learning comparisons. We tested the statistical significance of Z chromosome enrichment in this core gene list with an upper-tailed hypergeometric test, implemented in phyper, where each core gene is a sampling event without replacement from module assigned genes.

## Data and Code Availability

All raw data for this experiment is available on the NCBI Sequence Read Archive (accession: PRJNA698257). The count matrix, quality control results, and analysis code is available online (https://github.com/mattisabrat/sex-and-song).

---

**Figure S1 -.**
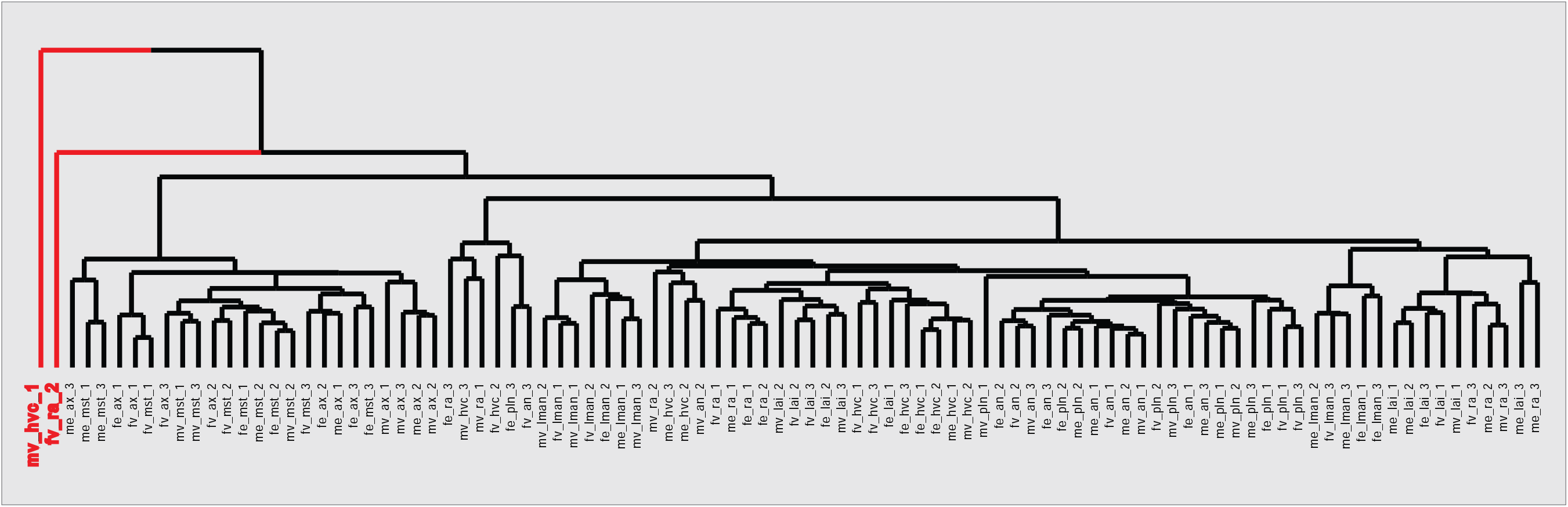
Outlier sample detection by hierarchical clustering. Two samples (a vehicle-treated male HVC sample and an E2-treated female RA sample, in red) form single sample branches in the hierarchical clustering tree, indicative of technical outliers unlikely to fit the correlational structure of the larger dataset. Samples were removed prior to gene network construction and module detection.

**Figure S2 -.**
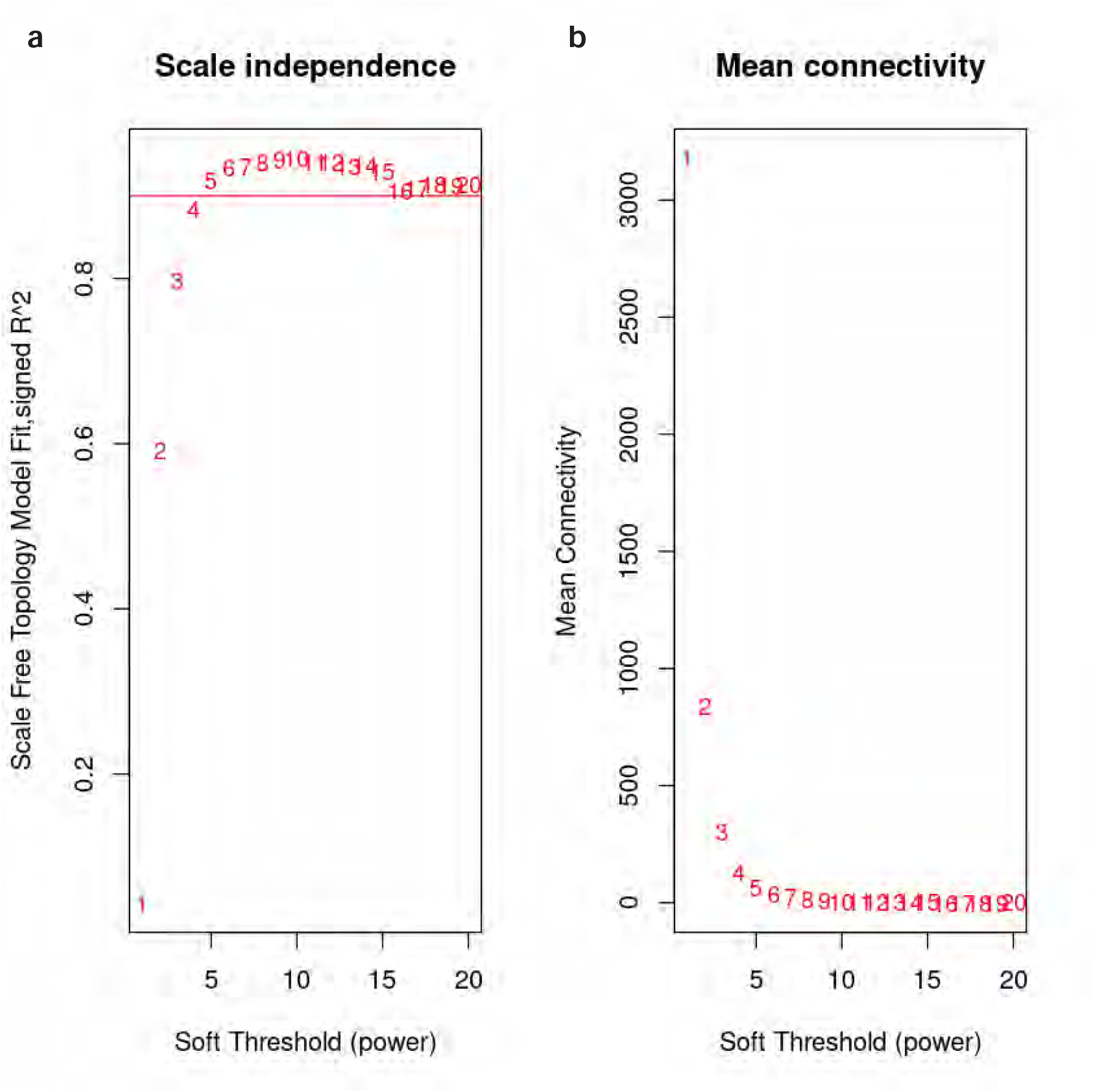
Selection of soft thresholding power for WGCNA model. Soft-thresholding power (beta, x-axis) is the exponent to which each element in the gene-to-gene correlation matrix is raised during adjacency matrix calculation. **a,** Scale-free fit index (y-axis) as a function of the soft-thresholding power (x-axis). Horizontal line indicates a fit of 90%. **b,** Mean connectivity (degree) in the network model (y-axis) as a function of the soft-thresholding power. We selected a power of 6 as it is on the knee of both plots and above the 90% scale free fit criteria.

**Figure S3 -.**
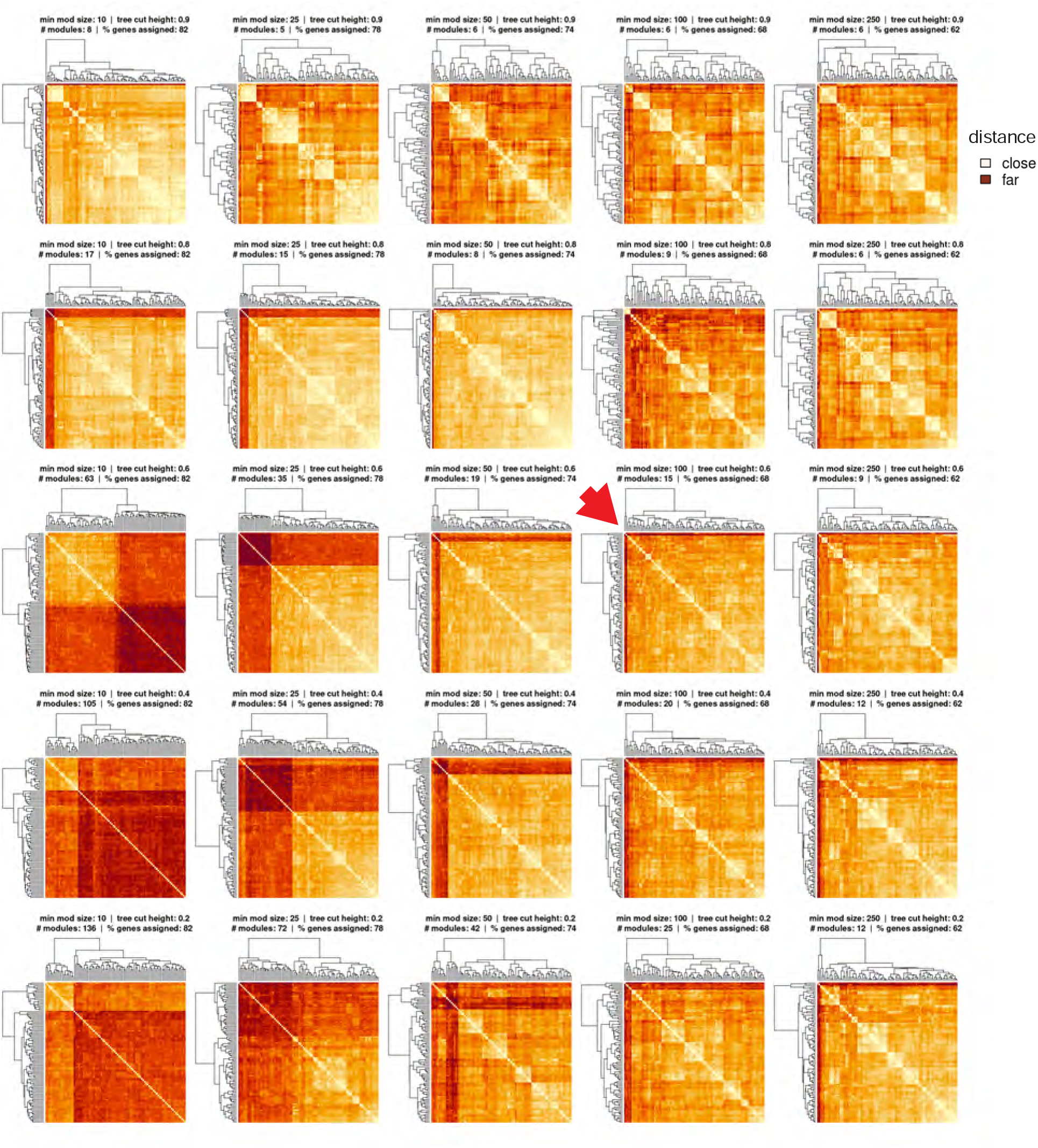
Selection of minimum module size and tree cut height parameter values for WGCNA model. Each plot shows the sample distance matrix that results from the parameters in the plot title. Titles also show the number of modules found in each resulting model and the percentage of genes in the finch genome assigned. Minimum module size increases across columns (left to right: 10, 25, 50, 100, 250) and tree cut height decreases down rows (top to bottom: 0.9, 0.8, 0.6, 0.4, 0.2). Red arrow indicates selected model. Model was selected to explain as much transcriptomic variance as possible while minimizing the number of technical- ly overfit samples.

**Figure S4 -.**
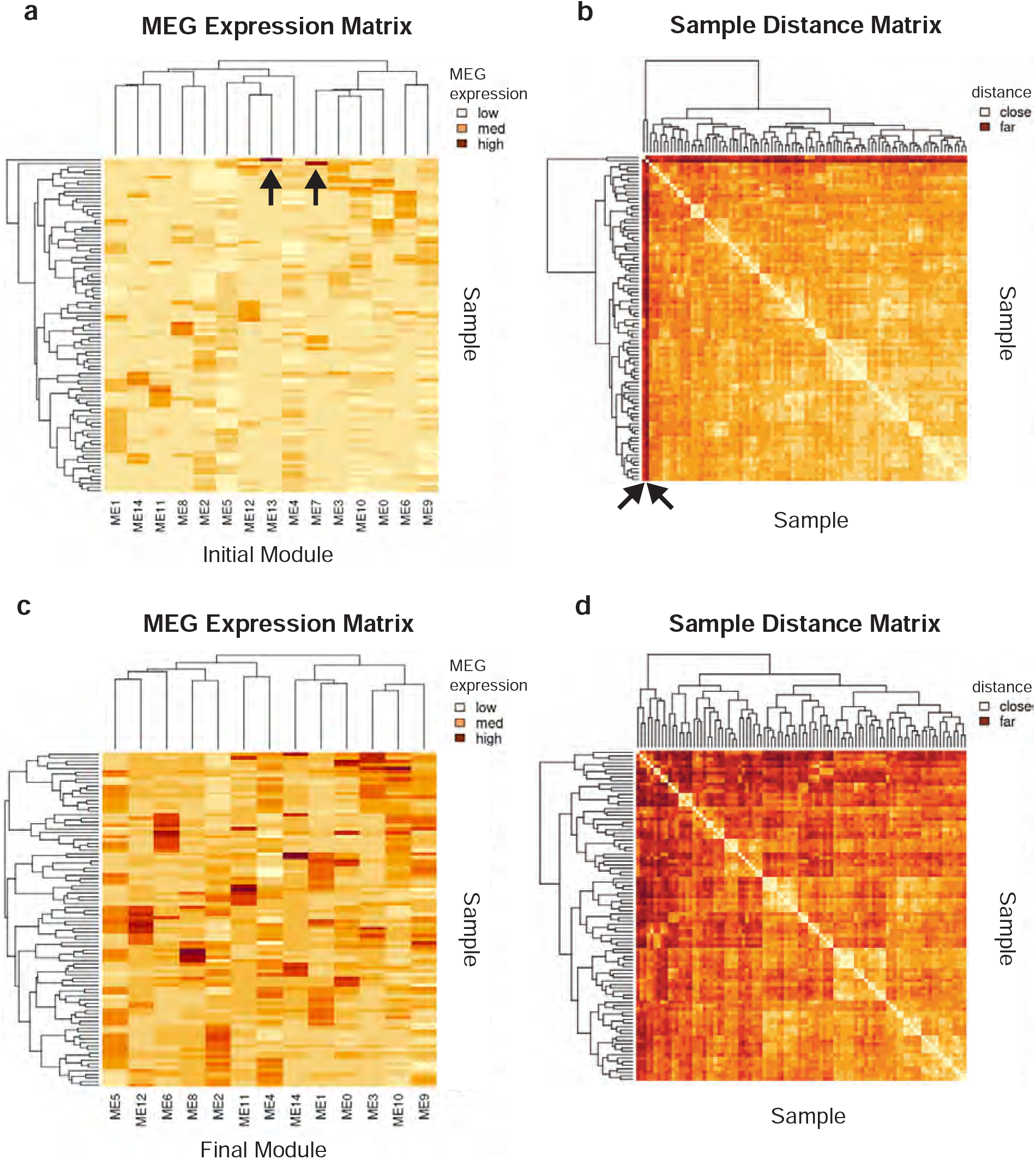
Initial module overfitting to single samples. **a,** Module eigengene (MEG) 7 and 13 are both highly expressed only in sinlge samples, indicated by black arrows. **b,** This overfitting causes these samples to be deep outliers in the sample-sample distance matrix, distant from all samples but tthemselves, indicated by black arrows. **c,** Removing these module eigengenes from the set prevents these samples from behaving as outliers in the distance matrix (**d**). These overfit modules were removed prior to module lettering and statistical analysis.

**Figure S5 -.**
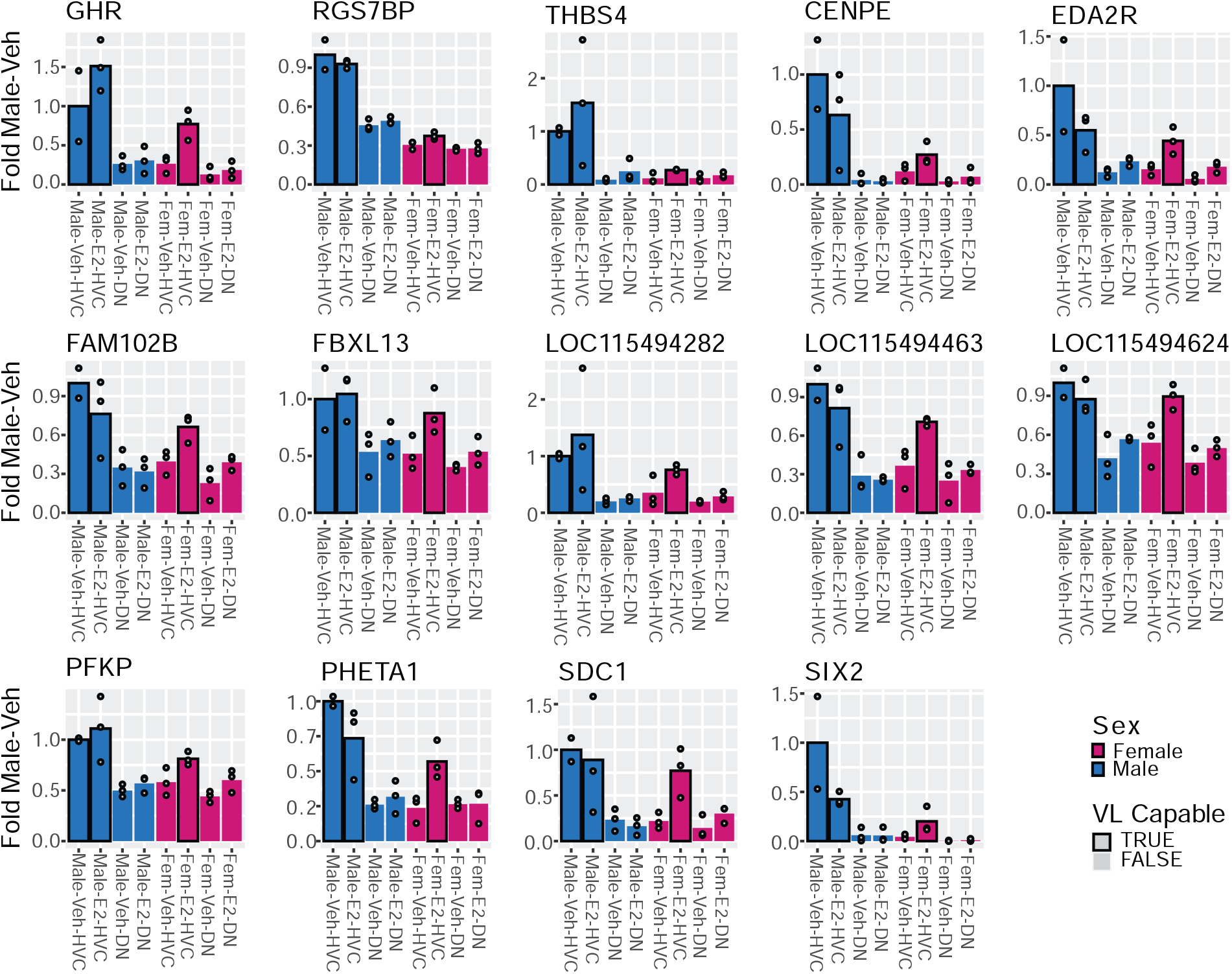
Expression of module G core genes in HVC and surrounding dorsal nidopallium. Each of the 14 core genes show reduced expression in female HVC relative to the male with an increase in expression in response to E2 treatment. Bar represents mean with individual data points shown. This transcriptional response to E2 is not seen in the surrounding DN.

**Table S1-.**
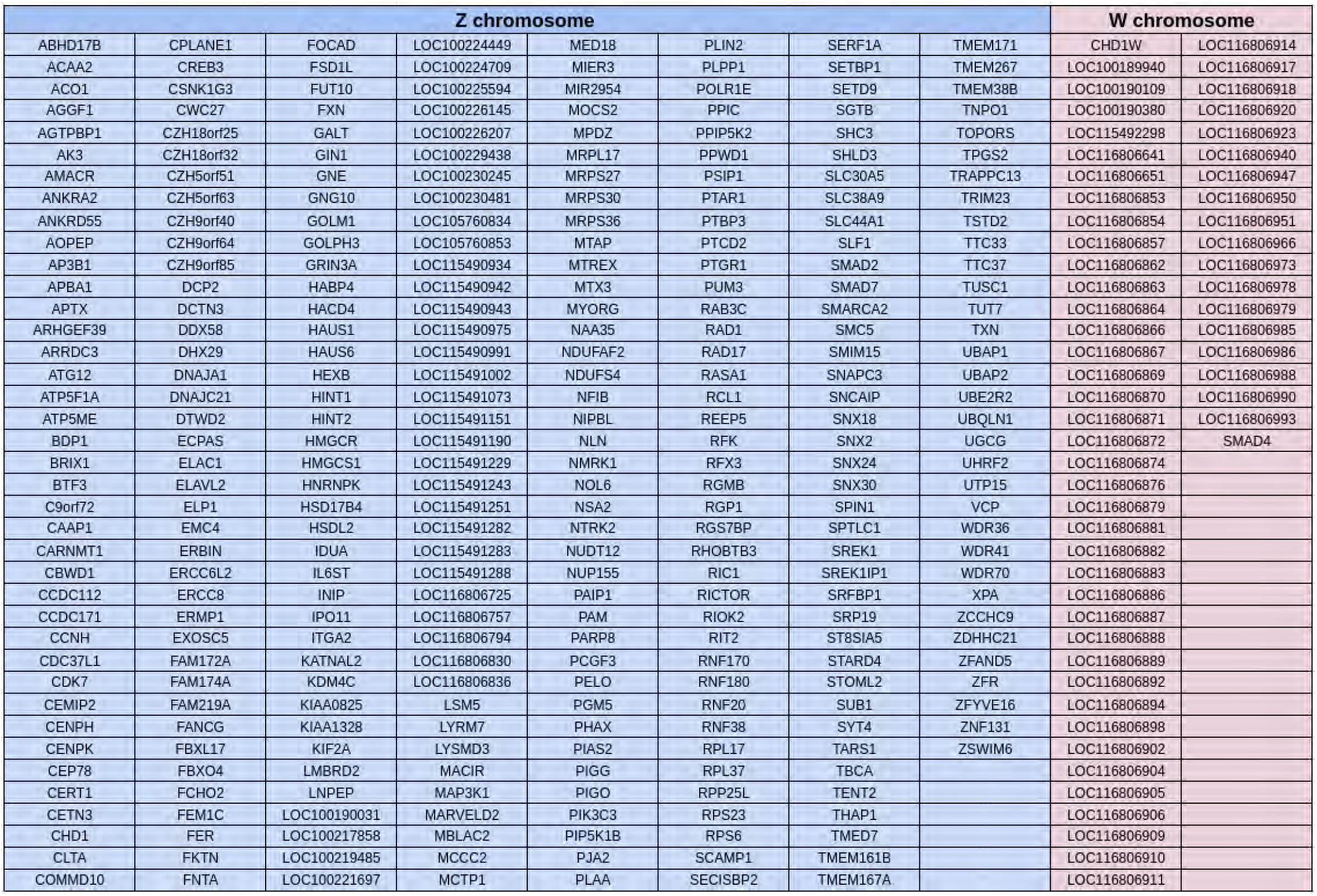
Sex chromosomes consistently repressed or expressed across regions. Lists Z and W chromosome genes from Fig. 3e-f which exhibited sexually dimorphic expression in vehicle-treated, non-vocal learning related samples across brain regions.

**Table S2 -.**
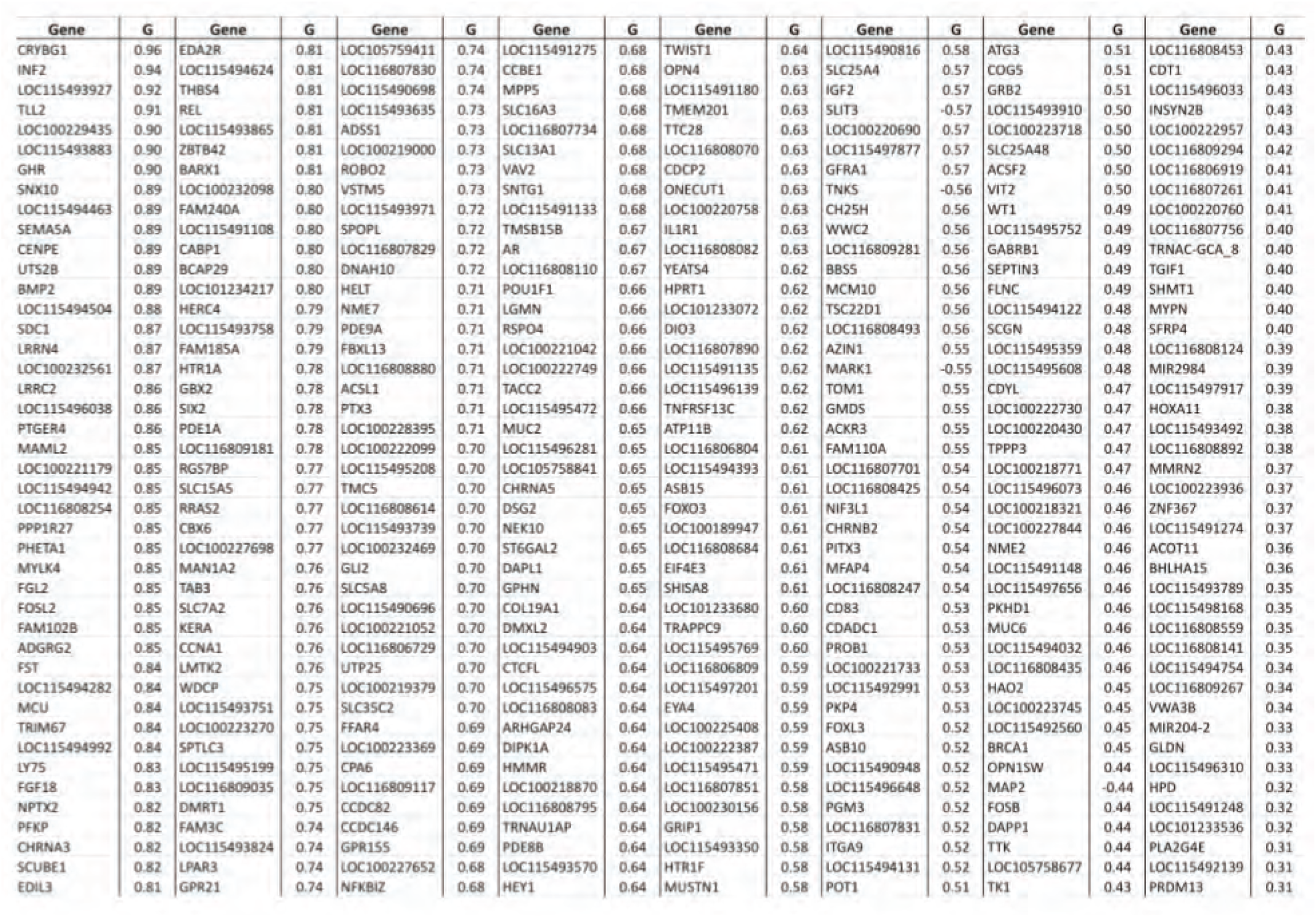
ModuleG constituent genes. Lists all genes assigned to module G by and their continuous membership in module G (Pearson r to MEG from moduleG)

## References

1. Jarvis, E. D. Evolution of vocal learning and spoken language. Science 366, 50–54 (2019).

2. Jarvis, E. D., Mirarab, S., Aberer, A. J., Li, B., Houde, P., Li, C., Ho, S. Y. W., Faircloth, B. C., Nabholz, B., Howard, J. T., Suh, A., Weber, C. C., da Fonseca, R. R., Li, J., Zhang, F., Li, H., Zhou, L., Narula, N., Liu, L., Ganapathy, G., Boussau, B., Bayzid, M. S., Zavidovych, V., Subramanian, S., Gabaldón, T., Capella-Gutiérrez, S., Huerta-Cepas, J., Rekepalli, B., Munch, K., Schierup, M., Lindow, B., Warren, W. C., Ray, D., Green, R. E., Bruford, M. W., Zhan, X., Dixon, A., Li, S., Li, N., Huang, Y., Derryberry, E. P., Bertelsen, M. F., Sheldon, F. H., Brumfield, R. T., Mello, C. V., Lovell, P. V., Wirthlin, M., Schneider, M. P. C., Prosdocimi, F., Samaniego, J. A., Vargas Velazquez, A. M., Alfaro-Núñez, A., Campos, P. F., Petersen, B., Sicheritz-Ponten, T., Pas, A., Bailey, T., Scofield, P., Bunce, M., Lambert, D. M., Zhou, Q., Perelman, P., Driskell, A. C., Shapiro, B., Xiong, Z., Zeng, Y., Liu, S., Li, Z., Liu, B., Wu, K., Xiao, J., Yinqi, X., Zheng, Q., Zhang, Y., Yang, H., Wang, J., Smeds, L., Rheindt, F. E., Braun, M., Fjeldsa, J., Orlando, L., Barker, F. K., Jønsson, K. A., Johnson, W., Koepfli, K.-P., O’Brien, S., Haussler, D., Ryder, O. A., Rahbek, C., Willerslev, E., Graves, G. R., Glenn, T. C., McCormack, J., Burt, D., Ellegren, H., Alström, P., Edwards, S. V., Stamatakis, A., Mindell, D. P., Cracraft, J., Braun, E. L., Warnow, T., Jun, W., Gilbert, M. T. P. & Zhang, G. Whole-genome analyses resolve early branches in the tree of life of modern birds. Science 346, 1320–1331 (2014).

3. Kumar, S. & Hedges, S. B. A molecular timescale for vertebrate evolution. Nature 392, 917–920 (1998).

4. van Tuinen, M. & Hadly, E. A. Error in estimation of rate and time inferred from the early amniote fossil record and avian molecular clocks. J. Mol. Evol. 59, 267–276 (2004).

5. Bolhuis, J. J., Okanoya, K. & Scharff, C. Twitter evolution: converging mechanisms in birdsong and human speech. Nat. Rev. Neurosci. 11, 747–759 (2010).

6. Doupe, A. J. & Kuhl, P. K. Birdsong and human speech: common themes and mechanisms. Annu. Rev. Neurosci. 22, 567–631 (1999).

7. Mooney, R. Neural mechanisms for learned birdsong. Learn. Mem. 16, 655–669 (2009).

8. Pfenning, A. R., Hara, E., Whitney, O., Rivas, M. V., Wang, R., Roulhac, P. L., Howard, J. T., Wirthlin, M., Lovell, P. V., Ganapathy, G., Mouncastle, J., Moseley, M. A., Thompson, J. W., Soderblom, E. J., Iriki, A., Kato, M., Gilbert, M. T. P., Zhang, G., Bakken, T., Bongaarts, A., Bernard, A., Lein, E., Mello, C. V., Hartemink, A. J. & Jarvis, E. D. Convergent transcriptional specializations in the brains of humans and song-learning birds. Science 346, 1256846 (2014).

9. Feenders, G., Liedvogel, M., Rivas, M., Zapka, M., Horita, H., Hara, E., Wada, K., Mouritsen, H. & Jarvis, E. D. Molecular mapping of movement-associated areas in the avian brain: a motor theory for vocal learning origin. PLoS One 3, e1768 (2008).

10. Gedman, G. L., Biegler, M. T., Haase, B., Wirthlin, M. E., Fedrigo, O., Pfenning, A. R. & Jarvis, E. D. Convergent gene expression highlights shared vocal motor microcircuitry in songbirds and humans. bioRxiv 2022.07.01.498177 (2022). doi:10.1101/2022.07.01.498177

11. Cahill, J. A., Armstrong, J., Deran, A., Khoury, C. J., Paten, B., Haussler, D. & Jarvis, E. D. Positive selection in noncoding genomic regions of vocal learning birds is associated with genes implicated in vocal learning and speech functions in humans. Genome Res. 31, 2035–2049 (2021).

12. Li, X., Wang, X.-J., Tannenhauser, J., Podell, S., Mukherjee, P., Hertel, M., Biane, J., Masuda, S., Nottebohm, F. & Gaasterland, T. Genomic resources for songbird research and their use in characterizing gene expression during brain development. Proc. Natl. Acad. Sci. U. S. A. 104, 6834–6839 (2007).

13. Lovell, P. V., Clayton, D. F., Replogle, K. L. & Mello, C. V. Birdsong ‘transcriptomics’: neurochemical specializations of the oscine song system. PLoS One 3, e3440 (2008).

14. Nottebohm, F. & Arnold, A. P. Sexual dimorphism in vocal control areas of the songbird brain. Science 194, 211–213 (1976).

15. Odom, K. J., Hall, M. L., Riebel, K., Omland, K. E. & Langmore, N. E. Female song is widespread and ancestral in songbirds. Nat. Commun. 5, 3379 (2014).

16. The zebra finch: a synthesis of field and laboratory studies. Choice 34, 34–5095–34–5095 (1997).

17. Bottjer, S. W., Glaessner, S. L. & Arnold, A. P. Ontogeny of brain nuclei controlling song learning and behavior in zebra finches. J. Neurosci. 5, 1556–1562 (1985).

18. 18. Garcia-Calero, E. & Scharff, C. Calbindin expression in developing striatum of zebra finches and its relation to the formation of area X. Journal of Comparative Neurology 521, 326–341 Preprint at 10.1002/cne.23174 (2013)

19. Konishi, M. & Akutagawa, E. Neuronal growth, atrophy and death in a sexually dimorphic song nucleus in the zebra finch brain. Nature 315, 145–147 (1985).

20. Nixdorf-Bergweiler, B. E. Divergent and parallel development in volume sizes of telencephalic song nuclei in male and female zebra finches. J. Comp. Neurol. 375, 445–456 (1996).

21. Nordeen, E. J. & Nordeen, K. W. Sex and regional differences in the incorporation of neurons born during song learning in zebra finches. J. Neurosci. 8, 2869–2874 (1988).

22. Shaughnessy, D. W., Hyson, R. L., Bertram, R., Wu, W. & Johnson, F. Female zebra finches do not sing yet share neural pathways necessary for singing in males. J. Comp. Neurol. 527, 843–855 (2019).

23. Holloway, C. C. & Clayton, D. F. Estrogen synthesis in the male brain triggers development of the avian song control pathway in vitro. Nat. Neurosci. 4, 170–175 (2001).

24. Mooney, R. & Rao, M. Waiting periods versus early innervation: the development of axonal connections in the zebra finch song system. J. Neurosci. 14, 6532–6543 (1994).

25. Gurney, M. E. & Konishi, M. Hormone-induced sexual differentiation of brain and behavior in zebra finches. Science 208, 1380–1383 (1980).

26. Gurney, M. E. Behavioral correlates of sexual differentiation in the zebra finch song system. Brain Research 231, 153–172 Preprint at 10.1016/0006-8993(82)90015-4 (1982)

27. Simpson, H. B. & Vicario, D. S. Early estrogen treatment alone causes female zebra finches to produce learned, male-like vocalizations. J. Neurobiol. 22, 755–776 (1991).

28. Simpson, H. B. & Vicario, D. S. Early estrogen treatment of female zebra finches masculinizes the brain pathway for learned vocalizations. J. Neurobiol. 22, 777–793 (1991).

29. Choe, H. N., Tewari, J., Zhu, K. W., Davenport, M., Matsunami, H. & Jarvis, E. D. Estrogen and sex-dependent loss of the vocal learning system in female zebra finches. Horm. Behav. 129, 104911 (2021).

30. Pohl-Apel, G. The correlation between the degree of brain masculinization and song quality in estradiol treated female zebra finches. Brain Res. 336, 381–383 (1985).

31. Pohl-Apel, G. & Sossinka, R. Hormonal determination of song capacity in females of the zebra finch: Critical phase of Treatment. Z. Tierpsychol. 64, 330–336 (1984).

32. Herrmann, K. & Arnold, A. P. Lesions of HVc block the developmental masculinizing effects of estradiol in the female zebra finch song system. J. Neurobiol. 22, 29–39 (1991).

33. Nordeen, K. W., Nordeen, E. J. & Arnold, A. P. Estrogen establishes sex differences in androgen accumulation in zebra finch brain. J. Neurosci. 6, 734–738 (1986).

34. Agate, R. J., Grisham, W., Wade, J., Mann, S., Wingfield, J., Schanen, C., Palotie, A. & Arnold, A. P. Neural, not gonadal, origin of brain sex differences in a gynandromorphic finch. Proc. Natl. Acad. Sci. U. S. A. 100, 4873–4878 (2003).

35. Gianaroli, L., Magli, M. C. & Ferraretti, A. P. in Brenner’s Encyclopedia of Genetics (Second Edition) (eds. Maloy, S. & Hughes, K.) 397–400 (Academic Press, 2013).

36. Konishi, M. & Akutagawa, E. A critical period for estrogen action on neurons of the song control system in the zebra finch. Proc. Natl. Acad. Sci. U. S. A. 85, 7006–7007 (1988).

37. Rhie, A., McCarthy, S. A., Fedrigo, O., Damas, J., Formenti, G., Koren, S., Uliano-Silva, M., Chow, W., Fungtammasan, A., Kim, J., Lee, C., Ko, B. J., Chaisson, M., Gedman, G. L., Cantin, L. J., Thibaud-Nissen, F., Haggerty, L., Bista, I., Smith, M., Haase, B., Mountcastle, J., Winkler, S., Paez, S., Howard, J., Vernes, S. C., Lama, T. M., Grutzner, F., Warren, W. C., Balakrishnan, C. N., Burt, D., George, J. M., Biegler, M. T., Iorns, D., Digby, A., Eason, D., Robertson, B., Edwards, T., Wilkinson, M., Turner, G., Meyer, A., Kautt, A. F., Franchini, P., Detrich, H. W., 3rd, Svardal, H., Wagner, M., Naylor, G. J. P., Pippel, M., Malinsky, M., Mooney, M., Simbirsky, M., Hannigan, B. T., Pesout, T., Houck, M., Misuraca, A., Kingan, S. B., Hall, R., Kronenberg, Z., Sović, I., Dunn, C., Ning, Z., Hastie, A., Lee, J., Selvaraj, S., Green, R. E., Putnam, N. H., Gut, I., Ghurye, J., Garrison, E., Sims, Y., Collins, J., Pelan, S., Torrance, J., Tracey, A., Wood, J., Dagnew, R. E., Guan, D., London, S. E., Clayton, D. F., Mello, C. V., Friedrich, S. R., Lovell, P. V., Osipova, E., Al-Ajli, F. O., Secomandi, S., Kim, H., Theofanopoulou, C., Hiller, M., Zhou, Y., Harris, R. S., Makova, K. D., Medvedev, P., Hoffman, J., Masterson, P., Clark, K., Martin, F., Howe, K., Flicek, P., Walenz, B. P., Kwak, W., Clawson, H., Diekhans, M., Nassar, L., Paten, B., Kraus, R. H. S., Crawford, A. J., Gilbert, M. T. P., Zhang, G., Venkatesh, B., Murphy, R. W., Koepfli, K.-P., Shapiro, B., Johnson, W. E., Di Palma, F., Marques-Bonet, T., Teeling, E. C., Warnow, T., Graves, J. M., Ryder, O. A., Haussler, D., O’Brien, S. J., Korlach, J., Lewin, H. A., Howe, K., Myers, E. W., Durbin, R., Phillippy, A. M. & Jarvis, E. D. Towards complete and error-free genome assemblies of all vertebrate species. Nature 592, 737–746 (2021).

38. Langfelder, P. & Horvath, S. WGCNA: an R package for weighted correlation network analysis. BMC Bioinformatics 9, 559 (2008).

39. Gedman, G., Haase, B., Durieux, G., Biegler, M. T., Fedrigo, O. & Jarvis, E. D. As above, so below: Whole transcriptome profiling demonstrates strong molecular similarities between avian dorsal and ventral pallial subdivisions. J. Comp. Neurol. (2021). doi:10.1002/cne.25159

40. Gedman, G. Songbird brain organization and its molecular convergence with humans for vocal imitation learning. (2021).

41. Lai, C. S., Fisher, S. E., Hurst, J. A., Vargha-Khadem, F. & Monaco, A. P. A forkhead-domain gene is mutated in a severe speech and language disorder. Nature 413, 519–523 (2001).

42. Teramitsu, I. & White, S. A. FoxP2 regulation during undirected singing in adult songbirds. J. Neurosci. 26, 7390–7394 (2006).

43. Kim, J., Lee, C., Ko, B. J., Yoo, D., Won, S., Phillippy, A., Fedrigo, O., Zhang, G., Howe, K., Wood, J., Durbin, R., Formenti, G., Brown, S., Cantin, L., Mello, C. V., Cho, S., Rhie, A., Kim, H. & Jarvis, E. D. False gene and chromosome losses affected by assembly and sequence errors. bioRxiv 2021.04.09.438906 (2021). doi:10.1101/2021.04.09.438906

44. Itoh, Y. & Arnold, A. P. Chromosomal polymorphism and comparative painting analysis in the zebra finch. Chromosome Res. 13, 47–56 (2005).

45. Warren, W. C., Clayton, D. F., Ellegren, H., Arnold, A. P., Hillier, L. W., Künstner, A., Searle, S., White, S., Vilella, A. J., Fairley, S., Heger, A., Kong, L., Ponting, C. P., Jarvis, E. D., Mello, C. V., Minx, P., Lovell, P., Velho, T. A. F., Ferris, M., Balakrishnan, C. N., Sinha, S., Blatti, C., London, S. E., Li, Y., Lin, Y.-C., George, J., Sweedler, J., Southey, B., Gunaratne, P., Watson, M., Nam, K., Backström, N., Smeds, L., Nabholz, B., Itoh, Y., Whitney, O., Pfenning, A. R., Howard, J., Völker, M., Skinner, B. M., Griffin, D. K., Ye, L., McLaren, W. M., Flicek, P., Quesada, V., Velasco, G., Lopez-Otin, C., Puente, X. S., Olender, T., Lancet, D., Smit, A. F. A., Hubley, R., Konkel, M. K., Walker, J. A., Batzer, M. A., Gu, W., Pollock, D. D., Chen, L., Cheng, Z., Eichler, E. E., Stapley, J., Slate, J., Ekblom, R., Birkhead, T., Burke, T., Burt, D., Scharff, C., Adam, I., Richard, H., Sultan, M., Soldatov, A., Lehrach, H., Edwards, S. V., Yang, S.-P., Li, X., Graves, T., Fulton, L., Nelson, J., Chinwalla, A., Hou, S., Mardis, E. R. & Wilson, R. K. The genome of a songbird. Nature 464, 757–762 (2010).

46. Itoh, Y., Melamed, E., Yang, X., Kampf, K., Wang, S., Yehya, N., Van Nas, A., Replogle, K., Band, M. R., Clayton, D. F., Schadt, E. E., Lusis, A. J. & Arnold, A. P. Dosage compensation is less effective in birds than in mammals. J. Biol. 6, 2 (2007).

47. Appenzeller, S., Schirmacher, A., Halfter, H., Bäumer, S., Pendziwiat, M., Timmerman, V., De Jonghe, P., Fekete, K., Stögbauer, F., Lüdemann, P., Hund, M., Quabius, E. S., Bernd Ringelstein, E. & Kuhlenbäumer, G. Autosomal-Dominant Striatal Degeneration Is Caused by a Mutation in the Phosphodiesterase 8B Gene. The American Journal of Human Genetics 86, 83–87 Preprint at 10.1016/j.ajhg.2009.12.003 (2010)

48. Kobarg, C. B., Kobarg, J., Crosara-Alberto, D. P., Theizen, T. H. & Franchini, K. G. MEF2C DNA-binding activity is inhibited through its interaction with the regulatory protein Ki-1/57. FEBS Lett. 579, 2615–2622 (2005).

49. James A. Cahill, Joel Armstrong, Alden Deran, Carolyn J. Khoury, Benedict Paten, David Haussler, Erich D. Jarvis. Positive selection in non-coding genomic regions of vocal learning birds is associated with genes implicated in vocal learning and speech functions in humans. Genome Res.

50. Chen, Y.-C., Kuo, H.-Y., Bornschein, U., Takahashi, H., Chen, S.-Y., Lu, K.-M., Yang, H.-Y., Chen, G.-M., Lin, J.-R., Lee, Y.-H., Chou, Y.-C., Cheng, S.-J., Chien, C.-T., Enard, W., Hevers, W., Pääbo, S., Graybiel, A. M. & Liu, F.-C. Foxp2 controls synaptic wiring of corticostriatal circuits and vocal communication by opposing Mef2c. Nat. Neurosci. 19, 1513–1522 (2016).

51. Girard, F., Eichenberger, S. & Celio, M. R. Thrombospondin 4 deficiency in mouse impairs neuronal migration in the early postnatal and adult brain. Mol. Cell. Neurosci. 61, 176–186 (2014).

52. Cáceres, M., Suwyn, C., Maddox, M., Thomas, J. W. & Preuss, T. M. Increased cortical expression of two synaptogenic thrombospondins in human brain evolution. Cereb. Cortex 17, 2312–2321 (2007).

53. Harris, S. E., Cox, S. R., Bell, S., Marioni, R. E., Prins, B. P., Pattie, A., Corley, J., Muñoz Maniega, S., Valdés Hernández, M., Morris, Z., John, S., Bronson, P. G., Tucker-Drob, E. M., Starr, J. M., Bastin, M. E., Wardlaw, J. M., Butterworth, A. S. & Deary, I. J. Neurology-related protein biomarkers are associated with cognitive ability and brain volume in older age. Nat. Commun. 11, 800 (2020).

54. Ruth, K. S., Day, F. R., Tyrrell, J., Thompson, D. J., Wood, A. R., Mahajan, A., Beaumont, R. N., Wittemans, L., Martin, S., Busch, A. S., Erzurumluoglu, A. M., Hollis, B., O’Mara, T. A., Endometrial Cancer Association Consortium, McCarthy, M. I., Langenberg, C., Easton, D. F., Wareham, N. J., Burgess, S., Murray, A., Ong, K. K., Frayling, T. M. & Perry, J. R. B. Using human genetics to understand the disease impacts of testosterone in men and women. Nat. Med. 26, 252–258 (2020).

55. Kornfeld, M., Synder, R. D., MacGee, J. & Appenzeller, O. The oculo-cerebral-renal syndrome of Lowe. Arch. Neurol. 32, 103–107 (1975).

56. Ates, K. M., Wang, T., Moreland, T., Veeranan-Karmegam, R., Ma, M., Jeter, C., Anand, P., Wenzel, W., Kim, H.-G., Wolfe, L. A., Stephen, J., Adams, D. R., Markello, T., Tifft, C. J., Settlage, R., Gahl, W. A., Gonsalvez, G. B., Malicdan, M. C., Flanagan-Steet, H. & Pan, Y. A. Deficiency in the endocytic adaptor proteins PHETA1/2 impairs renal and craniofacial development. Dis. Model. Mech. 13, (2020).

57. Garcez, R. C., Le Douarin, N. M. & Creuzet, S. E. Combinatorial activity of Six1-2-4 genes in cephalic neural crest cells controls craniofacial and brain development. Cell. Mol. Life Sci. 71, 2149–2164 (2014).

58. Gao, J., Kang, X.-Y., Sun, S., Li, L., Zhang, B.-L., Li, Y.-Q. & Gao, D.-S. Transcription factor Six2 mediates the protection of GDNF on 6-OHDA lesioned dopaminergic neurons by regulating Smurf1 expression. Cell Death Dis. 7, e2217 (2016).

59. Dehkhoda, F., Lee, C. M. M., Medina, J. & Brooks, A. J. The Growth Hormone Receptor: Mechanism of Receptor Activation, Cell Signaling, and Physiological Aspects. Front. Endocrinol. 9, 35 (2018).

60. Yuri, T., Kimball, R. T., Braun, E. L. & Braun, M. J. Duplication of accelerated evolution and growth hormone gene in passerine birds. Mol. Biol. Evol. 25, 352–361 (2008).

61. Rasband, S. A., Bolton, P. E., Fang, Q., Johnson, P. L. F. & Braun, M. J. Evolution of the Growth Hormone Gene Duplication in Passerine Birds. Genome Biol. Evol. 15, (2023).

62. Xie, F., London, S. E., Southey, B. R., Annangudi, S. P., Amare, A., Rodriguez-Zas, S. L., Clayton, D. F. & Sweedler, J. V. The zebra finch neuropeptidome: prediction, detection and expression. BMC Biol. 8, 28 (2010).

63. Frankl-Vilches, C. & Gahr, M. Androgen and estrogen sensitivity of bird song: a comparative view on gene regulatory levels. J. Comp. Physiol. A Neuroethol. Sens. Neural Behav. Physiol. 204, 113–126 (2018).

64. Friedrich, S. R., Nevue, A. A., Andrade, A. L. P., Velho, T. A. F. & Mello, C. V. Emergence of sex-specific transcriptomes in a sexually dimorphic brain nucleus. Cell Rep. 40, 111152 (2022).

65. Harvey, S., Baudet, M.-L. & Sanders, E. J. Growth hormone-induced neuroprotection in the neural retina during chick embryogenesis. Ann. N. Y. Acad. Sci. 1163, 414–416 (2009).

66. Phoenix, C. H., Goy, R. W., Gerall, A. A. & Young, W. C. Organizing action of prenatally administered testosterone propionate on the tissues mediating mating behavior in the female guinea pig. Endocrinology 65, 369–382 (1959).

67. Arnold, A. P. The organizational-activational hypothesis as the foundation for a unified theory of sexual differentiation of all mammalian tissues. Horm. Behav. 55, 570–578 (2009).

68. De Vries, G. J., Rissman, E. F., Simerly, R. B., Yang, L.-Y., Scordalakes, E. M., Auger, C. J., Swain, A., Lovell-Badge, R., Burgoyne, P. S. & Arnold, A. P. A model system for study of sex chromosome effects on sexually dimorphic neural and behavioral traits. J. Neurosci. 22, 9005–9014 (2002).

69. Chen, X., Grisham, W. & Arnold, A. P. X chromosome number causes sex differences in gene expression in adult mouse striatum. Eur. J. Neurosci. 29, 768–776 (2009).

70. McPhie-Lalmansingh, A. A., Tejada, L. D., Weaver, J. L. & Rissman, E. F. Sex chromosome complement affects social interactions in mice. Horm. Behav. 54, 565–570 (2008).

71. Cox, K. H. & Rissman, E. F. Sex differences in juvenile mouse social behavior are influenced by sex chromosomes and social context. Genes Brain Behav. 10, 465–472 (2011).

72. Bolger, A. M., Lohse, M. & Usadel, B. Trimmomatic: a flexible trimmer for Illumina sequence data. Bioinformatics 30, 2114–2120 (2014).

73. Dobin, A., Davis, C. A., Schlesinger, F., Drenkow, J., Zaleski, C., Jha, S., Batut, P., Chaisson, M. & Gingeras, T. R. STAR: ultrafast universal RNA-seq aligner. Bioinformatics 29, 15–21 (2013).

74. Liao, Y., Smyth, G. K. & Shi, W. featureCounts: an efficient general purpose program for assigning sequence reads to genomic features. Bioinformatics 30, 923–930 Preprint at 10.1093/bioinformatics/btt656 (2014)

75. Ewels, P., Magnusson, M., Lundin, S. & Käller, M. MultiQC: summarize analysis results for multiple tools and samples in a single report. Bioinformatics 32, 3047–3048 (2016).

76. Wickham, H., Averick, M., Bryan, J., Chang, W., McGowan, L., François, R., Grolemund, G., Hayes, A., Henry, L., Hester, J., Kuhn, M., Pedersen, T., Miller, E., Bache, S., Müller, K., Ooms, J., Robinson, D., Seidel, D., Spinu, V., Takahashi, K., Vaughan, D., Wilke, C., Woo, K. & Yutani, H. Welcome to the tidyverse. J. Open Source Softw. 4, 1686 (2019).

77. Wickham, H. ggplot2: Elegant Graphics for Data Analysis. (Springer Science & Business Media, 2009).

78. Luo, W., Friedman, M. S., Shedden, K., Hankenson, K. D. & Woolf, P. J. GAGE: generally applicable gene set enrichment for pathway analysis. BMC Bioinformatics 10, 161 (2009).

